# Single-cell analysis of myeloid cells in HPV^+^ tonsillar cancer

**DOI:** 10.1101/2022.10.04.510291

**Authors:** David Gomez Jimenez, Can Altunbulakli, Sabine Swoboda, Aastha Sobti, David Askmyr, Ashfaq Ali, Lennart Greiff, Malin Lindstedt

## Abstract

The incidence of Human Papillomavirus positive (HPV^+^) tonsillar cancer has been sharply rising during the last two decades. Myeloid cells represent an appropriate therapeutic target due to their ability to orchestrate antigen-specific immunity within the tonsil, the availability of viral antigens, and the proximity of the tumor and the underlying lymphoid tissue. However, the interrelationship of steady-state and inflammatory myeloid cell subsets, and their impact on patient survival remains unexplored. Here, we used single-cell RNA-sequencing to map the myeloid compartment in HPV^+^ tonsillar cancer. Our analysis unveiled the existence of four dendritic cell lineages, two macrophage polarization processes, and their sequential maturation profiles. We observed an expansion of the myeloid compartment in HPV^+^ tonsillar cancer, accompanied by interferon-induced cellular responses both in DCs and monocyte-macrophages. Within the DC lineages, we describe a balance shift in the frequency of progenitor and mature cDC favoring the cDC1 lineage in detriment of cDC2s, in HPV^+^ lesions. Furthermore, we observed that all DC lineages apart from DC5s matured into a common activated DC profile. In turn, the monocyte-macrophage lineage was subjected to early monocyte polarization events, which gave raise to inflammatory-activated, and chemokine-producing macrophages. We validated the existence of most of the single-cell RNA-seq clusters using 26-plex flow cytometry, and described a positive impact of cDC1, activated DCs and macrophages in patient survival using signature scoring. The current study contributes towards the understanding of myeloid ontogeny and dynamics in human papilloma driven tonsillar cancer, and details myeloid biomarkers that can be used to predict therapy effects and assess patient prognosis.

## INTRODUCTION

Tonsillar squamous cell carcinoma, or short tonsillar cancer (TC), is a head and neck cancer (HNC) caused by abnormal growth of epithelial cells of the tonsillar mucosa. Risk factors include tobacco smoking and alcohol abuse as well as high-risk Human Papillomavirus (HPV) infection (1). The incidence of HPV^+^ TC is sharply rising (2), and currently accounts for > 70% of TC cases (3, 4). Biomarkers associated with the immune compartment of the tumor microenvironment (TME) have shown prognostic value in HPV^+^ TC. For instance, several studies have highlighted a correlation between tumor-infiltrating CD8^+^ T-cells, T regulatory cells (Tregs) (5, 6), and PD-L1 expression (7, 8), respectively, to clinical outcome. However, immune-checkpoint blockade strategies, aiming at inducing T-cell responses, have shown limited clinical efficacy in HNCs, irrespective of HPV status (9, 10). A different approach to treat cancer concerns harnessing anti-tumor immunity by targeting myeloid cells due to their ability to modulate lymphoid cell activity (11). Increased knowledge about myeloid cell heterogeneity, their subset-specific features and functions within the TME of HPV^+^ TC, and their potential impact on survival is warranted in order to develop new treatment strategies.

Myeloid cells are divided into conventional dendritic cells (cDCs), monocytes, macrophages, and granulocytes, each characterized by cell-type specific functions (12, 13). DCs act as sentinel cells peripheral tissue by taking up antigens, transferring them into lymphoid organs, and triggering T-cell activation (12). In the context of HNC, both cDC1 and cDC2 have been shown to express CCR7 at protein level, highlighting their migratory potential (14). Furthermore, cDC2 subsets were specifically shown to induce and correlate with Th1 CD4^+^ T-cell abundance in the TME (14). In contrast, cDC1 have been described to migrate via XCL1 and CCL5, presumably produced by tumor infiltrating NK-cells (15). Using a combination of gene-signature scoring in bulk HNC RNA-seq, these chemokines were shown to correlate with both cDC1 and NK-cell signatures (15), stressing the relevance of the cDC1-NK-cell axis in HNC. Human cDC1s are known for their superior capacity to mount anti-tumoral CD8 T-cell responses through cross-presentation (16). However, *in vitro* studies have shown that human cDC2 from blood and tonsils also can cross-present antigens upon appropriate stimulation (16, 17). Compared to DCs, macrophages are sensors of tissue damage and infection, which help in clearing apoptotic cells and activating T-cells *in situ*. The functional role of macrophages in HNC has not been clarified, and several retrospective IHC studies have reported contradictory results regarding any prognostic impact (18–21). Studies of other cancers have presented macrophages as highly plastic cells involved in extracellular matrix remodeling through MMPs (22), angiogenesis via VEGF-α production (23), and attenuation of immune responses through TGF-β (24). Finally, monocytes can have both pro- and anti-tumoral functions, and differentiate into monocyte-derived DCs (25, 26) and tumor associated-macrophages (11, 27–29).

Recently, myeloid cell diversity has been revisited through the use of single-cell RNA-sequencing (scRNA-seq) in blood (30, 31), tonsil (32), and spleen (33) as well as in a variety of tumors (33–35), tumor draining lymph nodes (14, 36), and peritoneal ascites from cancer patients (26, 28). Although these studies report highly overlapping DC subsets, they also highlight differences in DC biology depending on the tissue of origin. In blood, the DC compartment, in addition to cDC1 and cDC2, also consists of the CD14^+^ CD163^+^ DC3 and Axl^+^ Siglec6^+^ DC5 subtypes (30). Analyses of the DC lineage in inflamed tonsil and spleen have demonstrated three additional populations closely related to cDC2s: a small proliferating DC cluster, a CLEC10A^−^CLEC4A^+^ cDC2 population, and an activated CCR7^+^ cDC2 subset (26, 33). The heterogeneity of the monocyte-macrophage lineage (Mono-Macs) is even larger, including up to 15 different communities in a recent cross-tissue meta-analysis (35). The diversity of the Mono-Mac lineage is partly related to the disparity of markers and clusters reported, but also due to the low resemblance of *in vivo* Mono-Macs with the well characterized *in vitro* M1/M2 macrophage models (37). Additionally, macrophage ontogeny has been postulated to condition Mono-Mac function, and tissue resident macrophage populations such as Langerhans cells (LC) have recently gained attention (36, 38, 39). Collectively, the divergences observed in scRNA-seq studies possibly relate to the differentiation and maturation of myeloid cells, which in tissue are greatly influenced by the local microenvironment.

Myeloid cells represent potential therapeutic targets in TC since they can orchestrate anti- and pro-tumor T-cell responses. Given their divergent T-cell polarization capacity (26, 31, 32, 40), there is a need to profile their diversity within the TME and healthy steady state settings. Several groups have assessed the transcriptional profiles of the whole immune compartment of HNC biopsies using scRNA-seq (41–43). However, these studies are limited in their capacity to resolve myeloid cell heterogeneity, due to low frequency of these cells in the TME. Furthermore, HNCs arise in different anatomical sites and are heterogeneous in terms of immune-infiltration, and thus, it is important to delineate tumor subsite-specific nuances. Lastly, published studies most often use blood or tonsil specimens from patients with local inflammation, which are informative, but may not represent appropriate steady state lymphoid tissue controls.

In this study, we evaluate myeloid cell heterogeneity and maturation status in HPV^+^ TC (*cf*. paired healthy tonsils (HT)), and the impact of these features on survival of HNC patients. We implement droplet-based 10X Genomics scRNA-seq to characterize FACS-sorted CD45^+^CD13^+^HLA-DR^+^ myeloid cells, describing transcriptomically unique clusters, which include novel cDC2, LC, and Mono-Mac populations. Notably, we describe a preferential recruitment of pre-cDC1s in the TME, giving rise to cDC1s that mature into activated DCs. We further show that patients enriched in gene-signatures from cDC1s and activated DCs and macrophages have a higher 5-year overall survival. Better understanding of the myeloid complexity and functionality in TC can improve the design of myeloid-targeted therapies, which potentially can overcome the limitations of T-cell oriented immunotherapies in TC.

## RESULTS

### Myeloid cells in HPV^+^ TC include subsets of DCs, monocytes and macrophages, and a small population of HLA-DR^+^ granulocytes

To evaluate the heterogeneity of the myeloid compartment in HPV^+^ TC, we dissociated fresh biopsies into single cells, and used flow cytometry to characterize and quantify CD45^+^CD13^+^HLA-DR^+^ myeloid cell populations (*SI Appendix*, Figure S1A). We observed a significant increase in the frequency of CD13^+^HLA-DR^+^ cells in TC (n=8) as compared to contralateral HT (n=5), indicating that the myeloid compartment expands in TC compared to HT (*SI Appendix*, Figure S1B). Within the myeloid gate, the greatest frequency increase in TC as compared to HT was observed in the Mono-Mac and LC populations, followed by cDC1s. Next, we addressed the diversity of myeloid cells in treatment naïve TC patients at transcriptomic level.

We sorted CD45^+^CD13^+^HLA-DR^+^ cells from five HPV^+^ TC biopsies and one paired contralateral HT, followed by scRNA-seq using the droplet-based 10X Genomics platform (Figure 1A). After QC, normalization, and sample integration, we used UMAP to visualize single cells in a low dimensional space. Unsupervised clustering of 9502 single cells revealed 12 clusters (Figure 1B; *SI Appendix*, Figure S1C), characterized by expression of MHC class II coding transcripts (e.g., *HLA-DRA, HLA-DRB1*) and heterogeneous expression of *CD14* (Figure 1C), with different distribution in TC and HT (Figure 1D).

**Figure 1.**
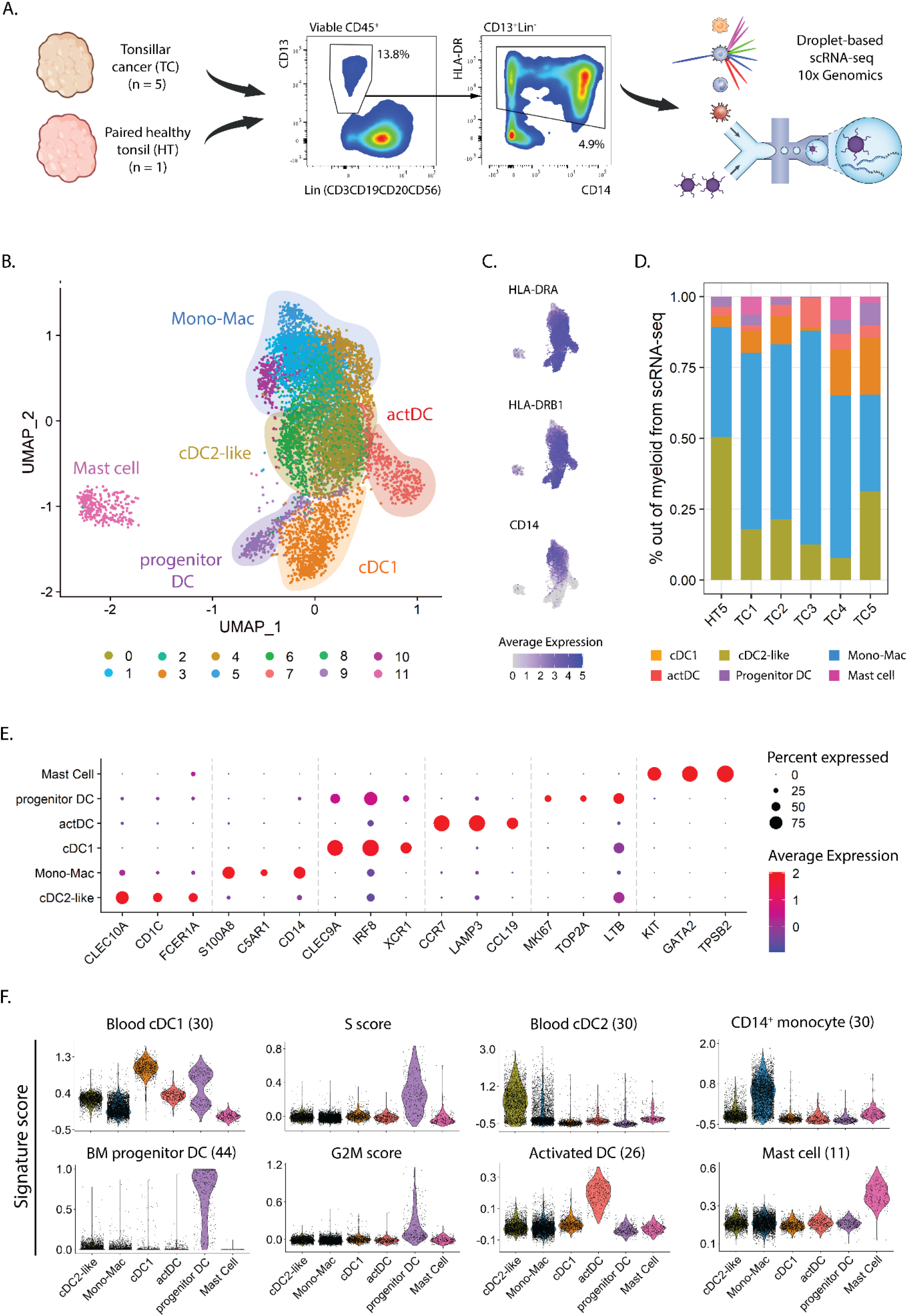
Heterogeneity of myeloid populations in TC and HT assessed by scRNAseq. **(A)** Viable CD45^+^ CD13^+^ HLA-DR^+^ myeloid cells were sorted from single cell suspensions of five TC samples and one contralateral HT and subjected to scRNA-seq using 10X genomics. The frequencies in the dot plots represent the average in TC calculated from viable CD45^+^ cells. **(B)** UMAP visualization of single cell transcriptomes displaying twelve clusters grouped into six major myeloid subtypes. **(C)** UMAP visualization, as in B, showing expression levels of *HLA-DRA, HLA-DRB1*, and *CD14*. **(D)** Frequency of myeloid populations across tissue of origin. **(E)** Dot plot displaying three canonical markers among the top ten differentially expressed genes across clusters. **(F)** Violin plots showing scores of indicated signatures. The score in each cell is shown in relative units, along with the density distribution shown by the shape of the plot. G2M and S scores represent the relative expression of cell cycle phase genes in a cell. Scale bars represent normalized counts in (C) and (E).

Next, we annotated the 12 clusters using a combination of canonical marker gene expression, after differential gene expression analysis (DGEA) across clusters and gene signature scoring (11, 26, 30, 44) (Figure 1E-F). The cDC1 population (cluster 3) expressed high levels of *CLEC9A, IRF8*, and *XCR1*, along with a high score of a described blood cDC1 gene-set. Similar to that observed at protein level (*SI Appendix*, Figure S1B), the cDC1 frequency in the scRNA-seq dataset increased in TC as compared to paired HT. In turn, cDC2-like cells (clusters 0 and 6) displayed *CLEC10A, CD1C*, and *FCER1A* expression and enrichment in a blood cDC2 gene set. Two additional DC clusters were predicted, representing different cell states of DC development and maturation. Progenitor DCs (cluster 9) exhibited markers related to cell cycle and progenitor cells (*MKI67, TOP2A*, and *LTB*), as illustrated by their high S and G2/M cell cycle phase scores. Additionally, this cluster featured high prediction scores of bone-marrow DC progenitor cells as well as low scores of hematopoietic stem cells and granulocyte-monocyte progenitors (*SI Appendix*, Figure S1D). The 4^th^ DC population, activated DC (actDC, cluster 7), was characterized by high expression of genes related to DC maturation, such as *CCR7, LAMP3* and *CCL19*, and high scores for a gene-set from activated DCs. Six clusters of CD14^+^ cells (clusters 1, 2, 4, 5, 8, and 10) conformed to the Mono-Mac lineage (Figure 1E-F). The Mono-Mac population was characterized by expression of canonical genes *S100A8/9* and *C5AR1*, as well as a high CD14^+^ monocyte signature score. As observed in the flow cytometry analysis (*SI Appendix*, Figure S1B), Mono-Macs were more frequent in TC as compared to HT (Figure 1D). Lastly, cluster 11 showed high expression of *KIT, GATA2*, and *TPSB2* transcripts as well as an exclusive mast cell signature score, which only was detected in TC samples. Mast cells expressed lower levels of MHC class II related transcripts compared to other myeloid cells, but expression of co-stimulatory molecule coding genes was not detected (Figure 1C; *SI Appendix*, Figure S1E).

### DC Lineage-specific subpopulations and analysis of cell identity dynamics

To detect rare populations, we divided the dataset into DCs and Mono-Macs, and re-clustered these lineages separately. Re-clustering followed by DGEA revealed 7 transcriptomically different communities in the DC lineages across TC and HT (Figure 2A-C; *SI Appendix*, Dataset S1). Three clusters of cDC2-like cells were characterized by expression of *CLEC10A, CD1C*, and *FCER1A* (Figure 2C), high scoring of blood and tissue cDC2 gene-sets (*SI Appendix*, Figure S2A), and heterogeneous distribution in TC and HT (Figure 2B). We identified two clusters of cDC2s with different expression of the transcriptional factor *RUNX3*. RUNX3^−^ cDC2s exhibited preferential expression of *CLEC10A, JAML, and TXNIP* and were only present in HT. In contrast, RUNX3^+^ cDC2s were characterized by expression of interferon inducible genes (*IFI30, IFITM3*), and were more abundant in TC as compared to HT. Additionally, MAFB^+^ LCs uniquely expressed *CD1A, CD14*, and *C1QA/B/C* and were only found in TC. This population was enriched in a LC and moDC gene-sets, but not in a previously described blood CD14^+^ DC3 gene-signature (*SI Appendix*, Figure S2A). MAFB^+^ LC displayed high G2M scores, highlighting their self-renewal capacity and further confirming their identity as LC (38) (*SI Appendix*, Fig S2C). A rare population of DC5s expressing *SIGLEC6, LILRA4*, and *IL3RA* was found closely associated with the cDC2-like cell clusters. DC5s featured a high blood DC5 gene-set score (*SI Appendix*, Figure S2A), and an increased frequency in HT as compared to TC. Lastly, cDC1s, actDCs, and progenitor DCs corresponded to the three remaining DCs populations, equally observed in the previous clustering (Figure 1B). The correspondence between the two clustering steps showed the homogeneity of cDC1, actDC, and progenitor DC communities at the present sequencing depth.

**Figure 2.**
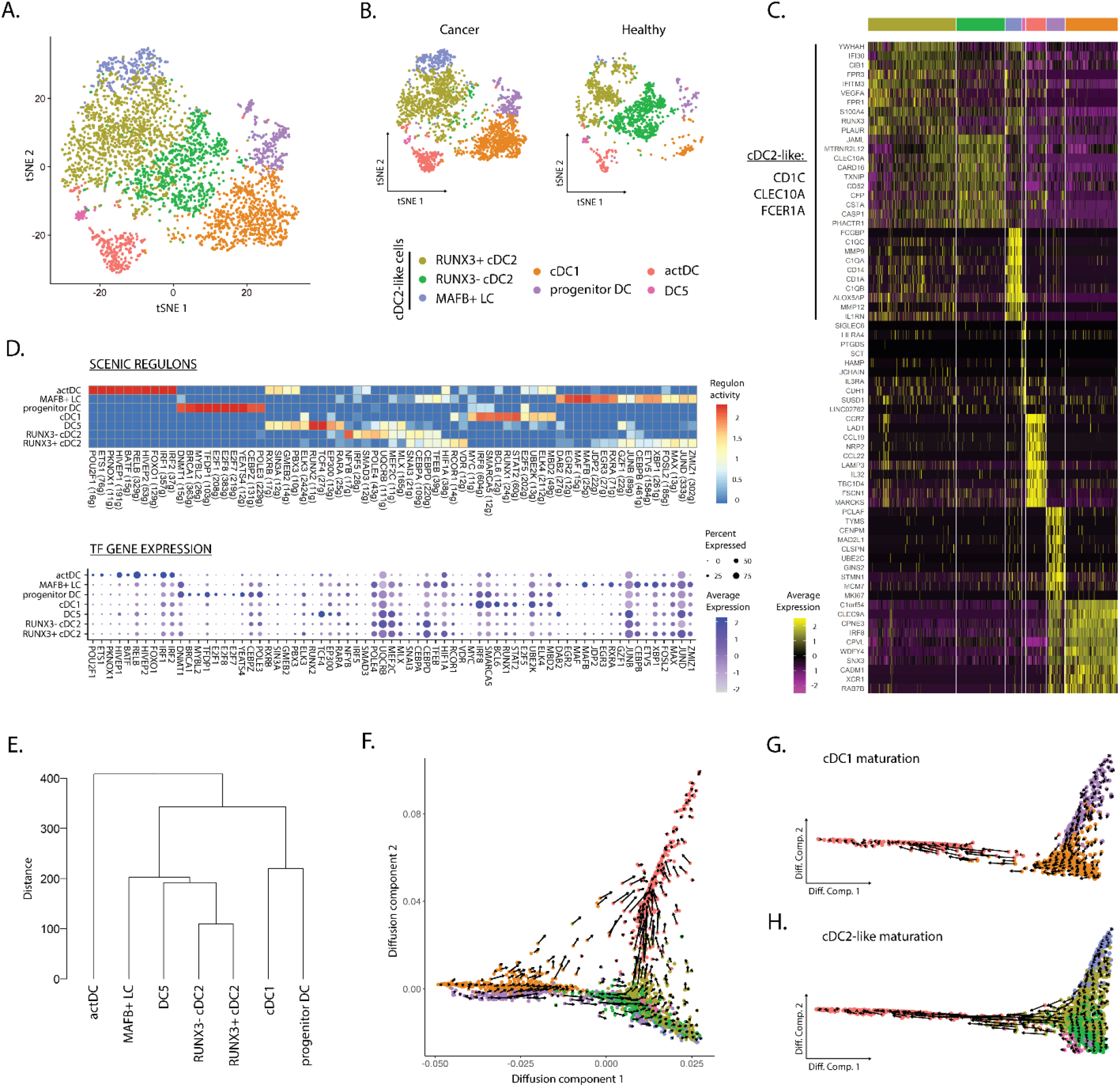
Dendritic cell dynamics and activation differ according to the tissue of origin. (**A**) tSNE plot showing seven distinct DC clusters independent of (**B**) or separated by tissue of origin. (**C**) Heatmap displaying top ten DEG per cluster after DGEA (FC > 1.5, p-value < 0.05). (**D**) Heatmap of the top 10 regulons per cluster colored by regulon activity (upper), and dot plot displaying the gene expression levels of each regulon’s TF per cluster (lower). (**E**) Dendrogram of DC clusters based on the Euclidean distance. Diffusion map combined with RNA velocity analysis to infer the development trajectory of all DCs (**D**), cDC1s (**E**) and DC2-like cells (**F**). DEG: differentially expressed genes; DGEA: differential gene expression analysis

We used SCENIC to assess different activity of gene regulatory networks in our clusters (Figure 2D; *SI Appendix*, Dataset S2). We noted that, cDC1s progenitor DCs and actDCs featured the most distinct regulons in the DC compartment, while cDC2-like cells and DC5s partly shared their regulatory networks. actDCs were characterized by high activity of RELB, HIVEP1 and IRF1/2 regulons. In turn, cDC1s featured high activity, and expression of STAT2, RUNX1, and IRF8 regulons. Progenitor DCs uniquely displayed activity and expression of E2F1/7/8 as well as BRCA1 and MYBL2. DC5s uniquely showed TCF4 and RUNX2 regulon activity and expression, and preferential RARA enrichment as compared to MAFB^+^ LC. On the other hand, MAFB^+^ LC exhibited exclusive MAFB activity and expression, and preferential enrichment in RXRA and DAB2 among other regulons and TFs. Finally, RUNX3^−^, RUNX3^+^ cDC2s, and MAFB^+^ LC shared several active regulons including SNAI3, CEBPD, and TFEB. However, while RUNX3^+^ cDC2s exhibited higher activity of VDR, RUNX3^−^ cDC2s were characterized by higher NFYB and SMAD3 activity. Next, we performed hierarchical clustering based on Euclidean distance between the clusters to assess their transcriptomic similarity (Figure 2E). We observed that actDCs were the most divergent group, clustering separately in the dendrogram. cDC2-like cells and DC5s clustered together and were distant from the cDC1 and progenitor DC pair, in line with the results from our gene regulatory network analysis.

Next, we sought to study cell identity dynamics of DC communities; thus, we projected single cell transcriptomes along with RNA velocity information onto diffusion map embeddings to infer future states of cells (Figure 2F-H). Overall, we observed directionality in RNA velocity of both cDC1 and cDC2-like cell branches towards actDCs. cDC1s and cDC2-like maturation into actDCs was confirmed by the sequential loss of *CLEC9A* and *CLEC10A*, respectively. These events were followed by a sequential increase in *LAMP3* expression along the trajectory, as observed in a 3D UMAP embedding (*SI Appendix*, movie S1). The maturation of cDC1 lineage was investigated by subdividing cDC1s along with progenitors enriched in cDC1 gene-set and actDC. RNA velocity analysis displayed a continuum of states along the progenitors into the fully differentiated cDC1 cluster, progressing to actDCs (Figure 2E). We evaluated transcriptomic changes along the developmental trajectories by assessing the velocity of highly variable genes with cluster-specific splicing dynamics (*SI Appendix*, Fig S2B). We observed that progenitor DCs had higher unspliced levels of DNAJB1, FUCA1 and DNMT1, on their development towards cDC1s. In turn, *SGO2* and *CTSL* were downregulated on the progression of progenitor DCs towards cDC1s and actDCs. cDC1s matured into actDCs displaying an initial upregulation of *CD40*, followed by *CCR7, PRDM1, NFKB1, LAMP3, IDO1*, and *MIR155HG*. In parallel, we analyzed cDC2-like cells likewise and detected a strong directionality from RUNX3^−^ cDC2s and MAFB^+^ LC into RUNX3^+^ cDC2s, which later progressed into actDCs (Figure 2H). We observed that all DC2-like clusters presented active transcription of *CD1C, IRF4*, and *HIVEP1*, and the last 2 genes were markedly upregulated upon maturation into actDCs (*SI Appendix*, Fig S2C). MAFB^+^ LC and RUNX3^+^ cDC2s exhibited upregulation of *RUNX3*, while only RUNX3^−^ cDC2s upregulated *ISG20* upon progression into RUNX3^+^ cDC2s. MAFB^+^ LCs specifically downregulated *CD1A* and *CD207* upon progression into RUNX3^+^ cDC2s. DC5s displayed active transcription of *TCF4*, while *LILRA4* was downregulated. In turn RUNX3^+^ cDC2s sequentially upregulated *CD274, LAMP3, TCF4*, followed by *IRF1*, upon maturation into actDCs. Contrary to progenitor cDC1s, we did not detect a directional flow in DC5s nor in CLEC10A^+^ progenitor DCs.

### Mono-Mac subsets display tumor-stage specific patterns

Similar to DC lineages, we subsetted Mono-Macs and re-clustered this lineage to address its heterogeneity. Using SLM clustering algorithm, followed by DGEA, we detected and characterized 6 transcriptomically unique clusters of Mono-Macs, with different distributions based on TNM classification of TCs (Figure 3A-C; SI *Appendix*, Figure S3A and Dataset S3). Monocytes, expressing high levels of *FCN1, CTSS*, and *JUNB*, were highly enriched in blood CD14^+^ and CD16^+^ monocyte signatures. In turn, ISG Monos featured high levels of IFN inducible genes (*IFIT1, IFIT3, IFITM*), and a high score of an ISG Mono gene-set. NLRP3 Macros expressed high levels of *NLRP3, CD300E*, and *AREG*, and were preferentially enriched in an NLRP3 gene-set. C1Q Macros expressed *C1QA/B/C* genes similarly to MAFB^+^ LC, but uniquely expressed *FCGR3A* and were enriched in a C1Q tumor-associated macrophage (TAM) gene-set. A cluster displaying expression of *IL7R, CCR7, MIR155HG*, and *CD274* was annotated as activated macrophages (act Macro). Finally, CXCL Macros showed high expression of *CXCL1/3/5, SPP1*, and *MMP1* as well as high score of an SPP1 TAM gene-set. Remarkably, signature scores corresponding to M1 and M2 activation states (37) did not distinguished the Mono-Mac clusters, indicating that the M1/M2 dichotomy did not resolve the differences among these (SI *Appendix*, Figure S3A). Apart from CXCL Macros, all Mono-Mac clusters were found in HT and TC, regardless of tumor stage. Instead, CXCL Macros were only detected in advanced primary tumors (n=2, T3N1M0, Figure 3B).

**Figure 3.**
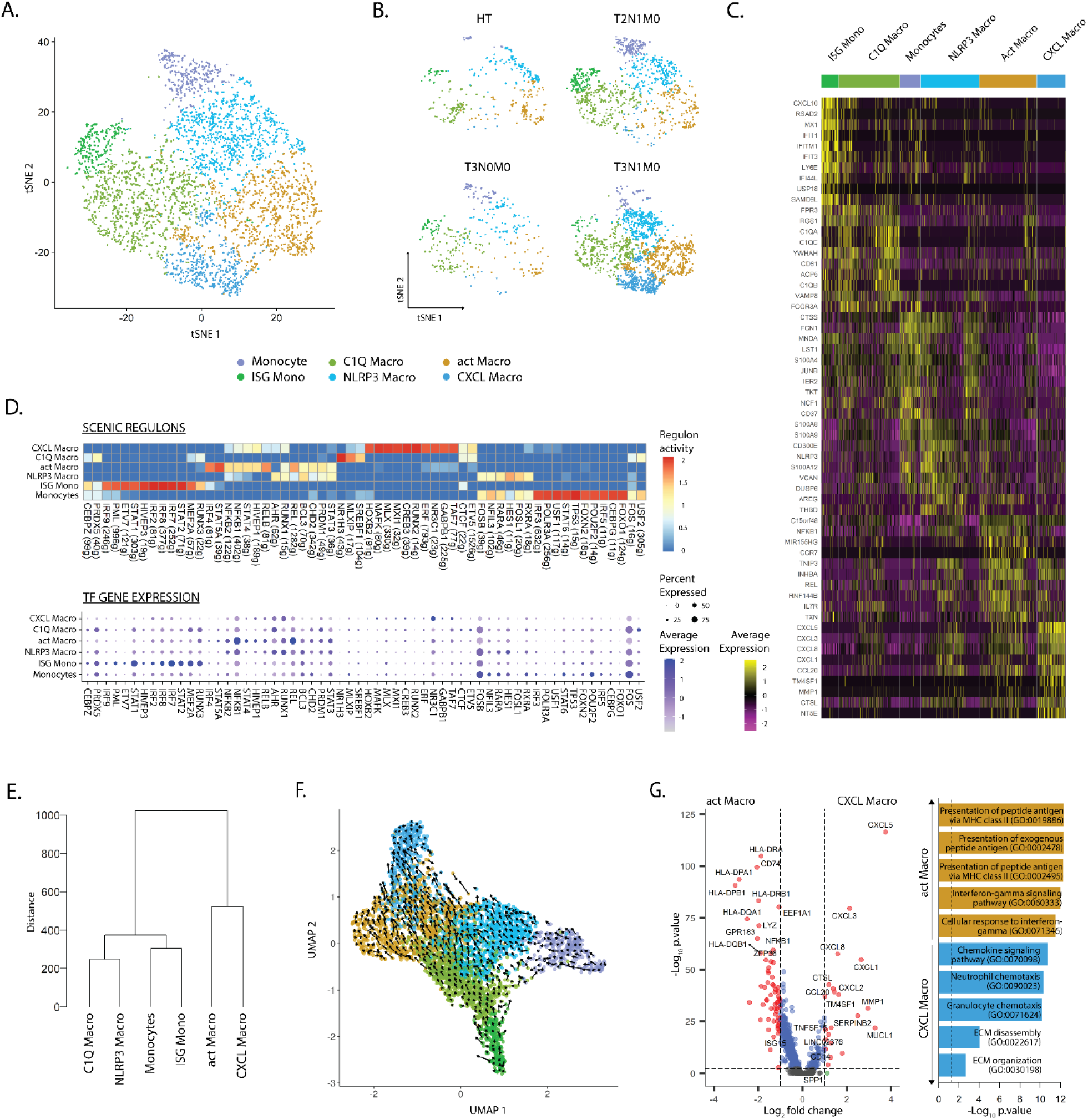
Mono-Mac subsets display tumor-stage specific patterns. (**A**) tSNE plot showing the six transcriptomically unique Mono-Mac clusters across cancer and healthy tissues. (**B**) tSNE plot of Mono-Mac distribution according to TNM classification. (**C**) Heatmap of the top 10 DEG per cluster after DGEA (FC > 1.5, p-value < 0.05). (**D**) Heatmap of the top 10 regulons per cluster colored by regulon activity (upper), and dot plot displaying the gene expression levels of each regulon’s TF per cluster (lower). (**E**) Dendogram of Mono-Mac clusters based on Euclidean distance. (**F**) UMAP plot showing the inferred development trajectory of Mono-Macs using RNA velocity. (**G**) DEG between CXCL and act Macro clusters, and enriched GO pathways obtained after GSEA.

We evaluated gene regulatory network activity and transcription factor expression in Mono-Mac clusters (Figure 3D; *SI Appendix*, Dataset S4). Briefly, the Mono-Mac lineage featured common expression of FOS and FOSB, where monocytes presented the higher regulon activity. In addition, Monocytes and NLRP3 Macros shared preferential expression and regulon activity of HES1, RARA, RXRA, and NFIL. In comparison, ISG Monocytes displayed expression and high regulon activity of IRF7/8 and STAT1, and shared RUNX3 and MEF2A regulon activity with C1Q Macros. Finally, CXCL and act Macros shared expression and activity of certain regulons including STAT4 and HIVP1, but while the former displayed preferential NR3C1 and TAF7 activity and expression, the latter featured higher STAT5A, NFKB1, and REL activity. We compared the transcriptomic similarity between the Mono-Mac clusters by performing hierarchical clustering based on Euclidean distance (Figure 3E). Monocytes and ISG monocytes, NLRP3 and C1Q Macros, and CXCL and act Macros, were paired together, respectively.

Next, we projected single cell transcriptomes along with RNA velocity information onto UMAP embeddings to decipher future state of cells (Figure 3F). We observed that Monocytes and ISG mono differentiated into NLRP3 and C1Q Macros, which in turn polarized into CXCL and act Macros, respectively. Furthermore, we evaluated transcriptomic changes across the Mono-Mac developmental trajectories by analyzing highly variable genes with dynamic changes in their velocity (SI *Appendix*, Figure S3B). We noted that Monocytes, ISG Monos, NLRP3, and C1Q Macros upregulated the expression of *IL1B* and *OLR1*, while the expression of these genes was drastically downregulated upon activation into CXCL and act Macros. Monocytes and NLRP3 Macros featured active transcription of *NLRP3, S100A9, AQP9* and *FCN1*. Upon progression into CXCL Macros these upregulated *MMP10* and *CXCL5*. On the contrary, ISG Monos presented active transcription of *ISG20* and *IDO1*, and sequentially upregulated *HLA-DPB1, IL4I1*, and *IRF4* as they differentiated into C1Q and act Macros, respectively. Together these results highlighted that there was a core transcriptional program in different stages of the Mono-Mac lineage trajectory.

To gain insights into the two parallel Mono-Mac differentiation processes, we performed pairwise DGEA between act and CXCL Macro clusters followed by GSEA (Figure 3G). act Macros featured a higher expression of antigen presentation related transcripts and enrichment of the corresponding pathways. In turn, CXCL Macros were characterized by high expression of chemotactic genes related to granulocyte and neutrophil chemotaxis (*CXCL1*/*3*/*5*/*8*), as well as genes related to matrix remodeling pathways (*MMP1, SPP1*). These results highlighted distinct functions of the terminal macrophage states in the two differentiation pathways present in our dataset.

### Validation of myeloid heterogeneity in TC and paired HT

Next, we sought to define a flow cytometry antibody panel to validate the myeloid communities detected by scRNA-seq at protein level. Protein targets were selected by performing DGEA analysis across myeloid cell clusters, followed by a filtering step to select cell membrane protein coding transcripts (SI *Appendix*, dataset S5). We evaluated TC and HT-derived single-cell suspensions using an antibody panel directed towards proteins encoded by genes identified in our scRNA-seq analysis (*XCR1, CD1c, CD14, CD5, CCR7, CD1A, CD207, SDC2, HLA-DRA, CD300E, FCGR3A*, and *TNFSF10*) (Figure 4A). The panel was complemented with antibodies specific for immune-checkpoints PD-L1, PD-L2, LAG3, and BTLA as well as the co-stimulatory protein CD40, given that their corresponding genes exhibited cluster/lineage specific expression in our scRNA-seq dataset (Figure 4B). Antibodies against CD19 and CD20 targeting B cells, and CD123 targeting plasmacytoid DCs, were included in the panel to increase resolution and compare myeloid cells to other APC populations. We also included other markers in our panel that, despite not being detected in our analysis, have recently been reported to mark novel myeloid populations. CD163 has been reported to mark a CD14^+^ DC3 population possibly distinct from CD14^+^ LC and moDC (30, 31). CCR2 was included in the panel since it is downregulated by monocytes upon differentiation (36, 45), while CD34 was included to detect cDC progenitors (30).

**Figure 4.**
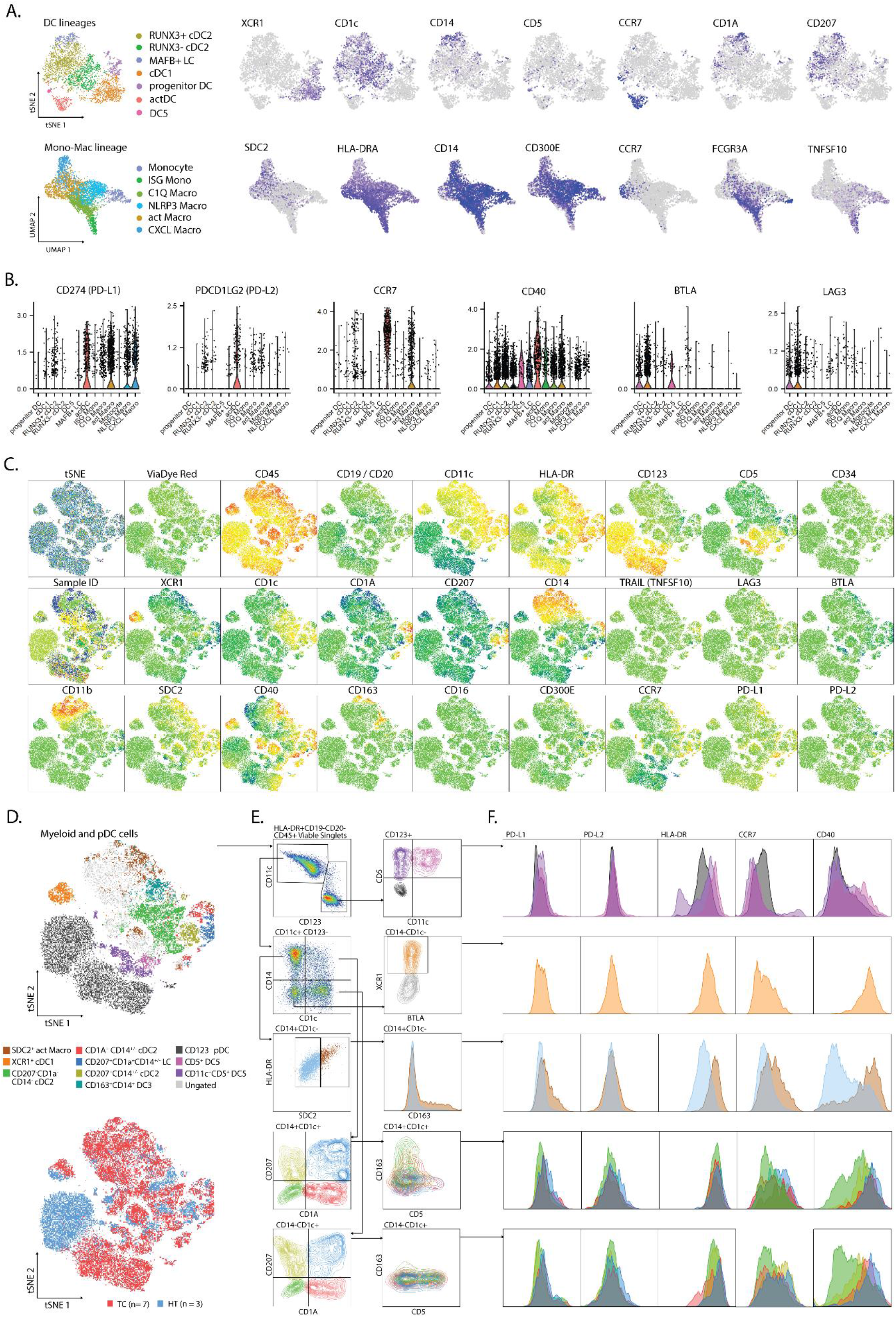
Validation of scRNA-seq clusters using 26-plex flow cytometry. (**A**) Gene expression of cluster-specific markers mapping to cell membrane, from scRNA-seq analysis. (**B**) Expression of genes related to APC function from scRNA-seq analysis. (**C**) tSNE plots showing individual marker expression on 45712 viable CD45^+^HLA-DR^+^CD19^−^CD20^−^ single cells from TC (n=7) and paired HT (n=3). (**D**) tSNE plots showing validated populations and their distribution in TC and HT tissues. (**E**) Gating strategy used to identify scRNA-seq clusters using flow cytometry. (**F**) Expression of genes related to APC function from flow cytometry.

We analyzed ten cryopreserved single cell suspensions, corresponding to biopsies from seven TCs and three paired HTs, by flow cytometry (*SI Appendix*, table S1). After singlet filtering, viable CD45^+^HLA-DR^+^CD19^−^CD20^−^ cells were concatenated and visualized within a tSNE space. We observed a clear signal from all markers apart from CD16, LAG3, TRAIL, BTLA, CD34, and CD300E (Figure 4C). We established a gating strategy to define several myeloid populations in human TC and HT and compare their levels of PD-L1, PD-L2, HLA-DR, CCR7, and CD40 (Figure 4D-F). We obtained a clear separation of the dataset based on CD11c and CD123, with minimal overlap between the two markers. Careful inspection of the CD123^+^ gate revealed two small CD5^+^ populations corresponding to DC5s, which were distinguishable by CD11c expression as previously reported in blood (30). Compared to pDCs, DC5s featured higher levels of HLA-DR, CD40, and CCR7. In turn, inspection of the CD11c^+^CD14^+^CD1c^−^ Mono-Mac gate revealed a parallel increase of HLA-DR along the SDC2 signal. SDC2^+^ HLA-DR^high^ corresponded to act Macros according to their co-expression of PD-L1, CD40, and CCR7 at protein level, as observed at RNA level. In addition, this population exclusively expressed CD163 compared to other CD14^+^ Mono-Mac populations. Next, we focused on resolving the heterogeneity of the CD1c^+^ gates. Similar to our scRNA-seq analysis, we observed co-expression of CD1c and CD207 in CD14^+/−^ cDC2s, suggesting that these corresponded to our newly identified RUNX3+ cDC2 cluster. In turn, CD1a^+^CD207^+^CD14^+/−^ were identified as LC. Interestingly, we observed the co-expression of CD163 and CD14 at protein level in CD1c^+^CD1A^−^CD207^−^ cells. This indicates that DC3s are present in TC as previously reported in blood (30, 31) and other HNCs (40), but are indistinguishable from other CD1c^+^ cells by scRNA-seq at our sequencing depth. In turn, CD5 expression inversely correlated with CD14 in CD1c^+^ cells, suggesting that these were “bona fide” cDC2s as described in blood (31). While all CD1c^+^ populations had some expression of PD-L1, PD-L2, HLA-DR, CCR7, and CD40, CD207^−^CD1A^−^CD1c^+^ cDC2s showed the most immature profile according to CD40 and CCR7 levels. Finally, XCR1^+^ cDC1s were clearly distinguishable by their high CD40 and HLA-DR levels, CCR7 expression, and residual PD-L1 and PD-L2 expression, indicating a mature profile of these population.

To assess transcriptomic changes induced by TME we performed DGEA comparing TC and HT myeloid populations from our scRNA-seq dataset (*SI Appendix*, Figure S4, Dataset S6). Progenitor DC in TC expressed higher levels of cDC1 canonical genes (*CLEC9A, BATF3, XCR1*) and cell cycle genes (*CCND1, CAMK2D*), as illustrated by the enrichment in blood cDC1 and cell cycle scores respectively. In contrast, progenitor DCs in HT featured higher levels of canonical cDC2 genes (*CLEC10A, CD1C*) and blood cDC2 gene-set score. On comparing DEG between TC and HT, we observed a common increase of IFN response-related genes (*ISG15, ISG20, TXN)* in all DC clusters. Conversely, all myeloid populations in TC showed a downregulation of inflammasome-regulation genes *MTRNR2L12, DNASE1L3*, and *LGALS2*, as well as MHC-II coding genes.

### cDC1s and activated DCs and macrophages correlate with increased survival in HNC patients

We sought to assess the presence of myeloid populations described here in TC and HT, in other HNCs. To this end, we accessed publicly available scRNA-seq data (41), subset myeloid cells and integrated them using our inhouse dataset as reference. We observed that all clusters identified in our inhouse TC dataset were also present in other subtypes of HNC, although with vxsarying frequencies (*SI Appendix*, Figure S5A). RUNX3^−^ cDC2s were only marginally present in HNC subtypes (4 predicted cells), indicating that this cell state was specific to healthy tissue. Surprisingly, the diversity of tumor-infiltrating myeloid cells was similar between HPV^+^ and HPV^−^ tumors (*SI Appendix*, Figure S5B).

To investigate the prognostic impact of myeloid populations in HNC, we defined gene signatures that enabled us to infer relative abundance of myeloid populations within bulk HNC RNA-seq. We performed DGEA across all myeloid clusters that were isolated from TC, obtaining a total of 998 DEG (FC ≥ 2, p-val ≤ 0.01, *SI Appendix*, dataset S5). Then, we filtered our candidate genes using a publicly available scRNA-seq HNC dataset (41), to select cluster-specific genes that were not expressed by other tumor infiltrating-leukocytes (Figure 5A-B). We obtained 3-gene transcriptomic signatures for all our clusters except progenitor DCs, monocytes, and C1Q Macros. We further investigated the impact of these 3-gene signatures within bulk RNA-seq transcriptomes of HNC from the TCGA (46). We applied cox proportional-hazards regression to compare the survival rate of high (Q4) and low (Q1) scoring quartiles of each gene signature (Figure 5C, Table 1). We found that high scores of cDC1, actDC, and act Macro gene-sets correlated with increased 5-year survival in HNC patients, independently of age and sex. Of note, high score of the cDC1 signature was also positively correlated to survival when TC patients were assessed separately (n=41, p.value = 0.024).

**Table 1.** Adjusted hazard ratios (HR) comparing the five-year overall survival rate between HNC patients with high (Q4) and low (Q1) cluster-specific gene-set scores. The multivariate HR model was corrected by gender and age at diagnosis. HR: hazard-ratio; CI: confidence interval; N: number of patients. Statistically significant gene-signatures are indicated in bold.

| CELL-SPECIFIC<br>GENE<br>SIGNATURE | UNIVARIATE MODEL |  |  | MULTIVARIATE MODEL |  |  |
| --- | --- | --- | --- | --- | --- | --- |
|  | HR (95% CI) | p-value | N | HR (95% CI) | p-value | N |
| **cDC1** | **0.48 (0.3-0.77)** | **0.0023** | **225** | **0.47 (0.29-0.76)** | **0.0022** | **225** |
| cDC2-like | 0.94 (0.63-1.39) | 0.767 | 229 | 0.93 (0.61-1.4) | 0.738 | 229 |
| LC | 0.81 (0.54-1.21) | 0.316 | 226 | 0.93 (0.61-1.41) | 0.746 | 226 |
| **actDC** | **0.51(0.33-0.78)** | **0.0023** | **227** | **0.52 (0.34-0.81)** | **0.0036** | **227** |
| DC5 | 0.94 (0.61-1.44) | 0.788 | 222 | 0.93 (0.61-1.43) | 0.767 | 222 |
| ISG Mono | 0.85 (0.56-1.28) | 0.453 | 224 | 0.77 (0.5-1.17) | 0.234 | 224 |
| CXCL Macro | 1.19 (0.79-1.79) | 0.38 | 226 | 1.13 (0.75-1.7) | 0.557 | 226 |
| **act Macro** | 0.68 (0.45-1.03) | 0.074 | 225 | **0.64 (0.42-0.95)** | **0.042** | **225** |
| NLRP3 Macro | 1.01 (0.67-1.5) | 0.96 | 229 | 0.97 (0.65-1.45) | 0.894 | 229 |

**Figure 5.**
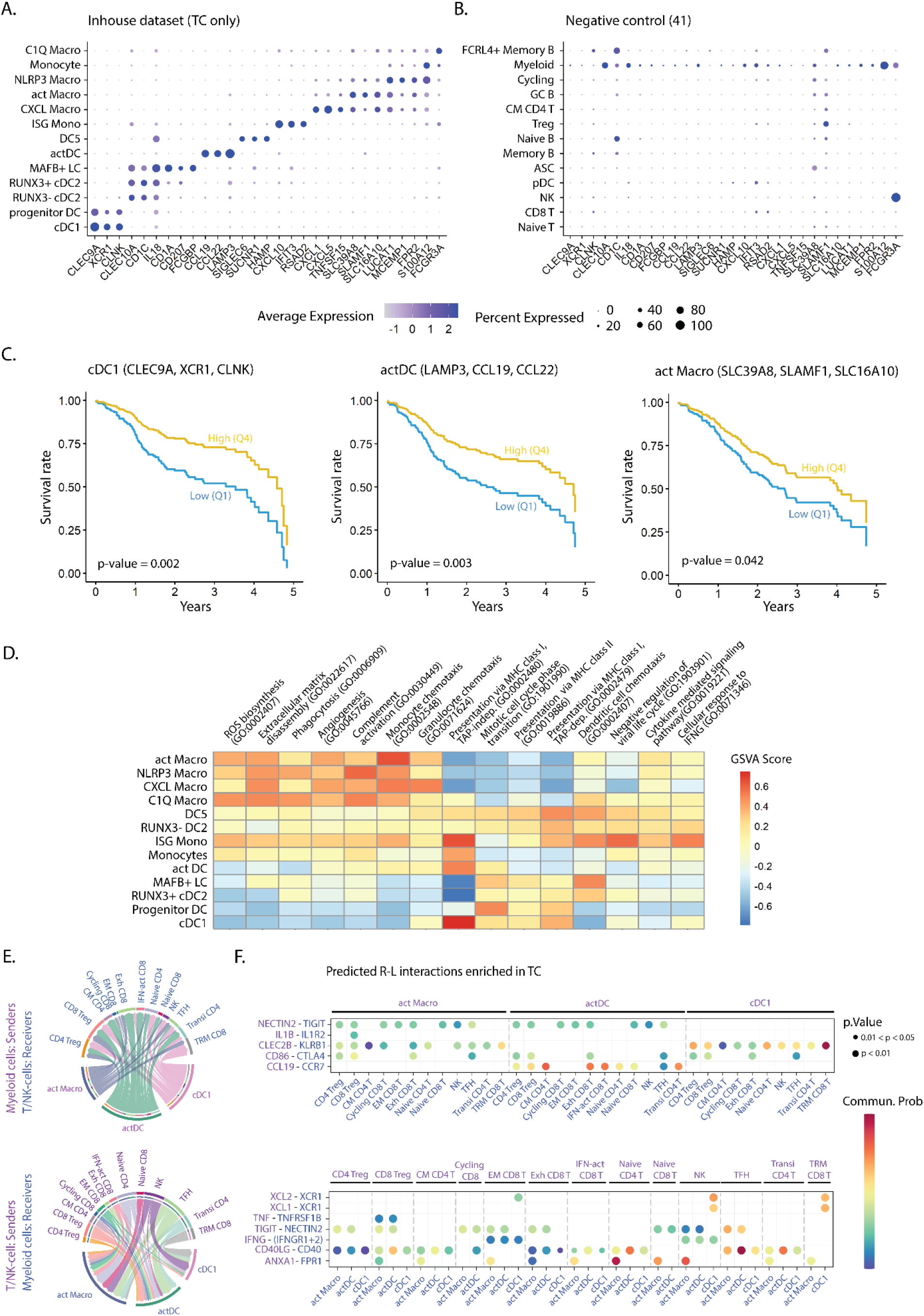
(**A**) Dot plot displaying cluster-specific signature genes in the myeloid dataset and (**B**) in a publicly available scRNA-seq dataset (41). **(C)** Survival curves showing 5-year overall survival rates for the indicated gene-sets, from a publicly available bulk RNA-seq dataset of HNC patients (n=530) (46). Comparison of survival probability was performed using log-rank test and cox-proportional hazards model. Years: years since diagnosis. (**D**) Heatmap colored by GSVA score of enriched GO pathways in the myeloid clusters (adjusted p-value < 0.05). (**E**) Chord diagrams displaying the interaction between cell pairs. Interaction strength is presented as the weight of the edges, and the receiver cell cluster is indicated by the apex of the arrow. (**F**) Bubble plot showing statistically significant receptor-ligand (R-L) coding gene pairs in TC between myeloid cell subsets and T/NK-cells from a publicly available dataset (41). DEG: differentially expressed genes; DGEA: differential gene expression analysis; GSVA: gene set variation analysis; FC: fold change; p: p-value; CM: central memory; EM: effector memory; Exh: exhausted; IFN-act: IFN-activated; NK: natural killer; TFH: T follicular helper; Transi: transitional; TRM: tissue-resident memory.

Next, we aimed at elucidating mechanisms by which cDC1s, actDCs, and act Macros impacted survival. Pathway enrichment analysis (GO bioprocesses 2018 (47), adjusted p-value < 0.05) revealed subset specific functions within the myeloid compartment in TC (*SI Appendix*, dataset S7). Highly scoring pathways related to myeloid cell function, such as antigen-presentation and chemotaxis, were subjected to GSVA score transformation and compared across clusters (Figure 5D). Overall, we found that all DCs, but also ISG monos, featured higher scores for antigen presentation pathways compared to macrophages. Specifically, the cDC1 cluster displayed the highest score in pathways related to cross-presentation via MHC-I, followed by ISG monos, DC5s, and actDCs. In contrast, all DC clusters presented similar scores in presentation via MHC-II, which was higher compared to Mono-Mac clusters. In turn, macrophage clusters were characterized by higher expression of genes related to chemotaxis, phagocytosis, ECM disassembly, angiogenesis, and complement pathway. Notably, ISG monocytes were preferentially enriched in cellular response to IFNG and negative regulation of viral life cycle, suggesting a TME specific polarization of this cluster.

To investigate which leukocytes interacted with cDC1s, actDCs and act Macros, we accessed a publicly available HNC scRNA-seq dataset (41). After annotating single cells from TC and HT based on canonical marker expression, we integrated all leukocytes of the external dataset and our inhouse dataset (*SI Appendix*, Figure S5C). Following normalization, we performed receptor ligand analysis to identify major cell-cell interactions in TC, as well as receptor-ligand gene pairs overexpressed in TC as compared to HT (*SI Appendix*, dataset S8). We focused our analysis on the interaction of myeloid cells with T/NK cell subsets, because of the high abundance of the last in the TME (Figure 5E-F). cDC1s featured higher signaling strength with NK and tissue resident memory (TRM) CD8 T-cells, followed by Treg subsets, as compared to other T/NK cells. Furthermore, cDC1s interacted with TRM CD8 T-cells, effector-memory (EM) CD8 T-cells, and NK cells via XCL1/XCL2-XCR1 as well as CLEC2B-KLRB1 axes. Based on the predicted signaling strength, actDCs and act Macros interactions were more diverse than cDC1s. Both actDCs and act Macros were predicted to interact with several T/NK subsets via NECTIN2-TIGIT. However, while actDCs were uniquely predicted to interact with T/NK subsets via CCL19-CCR7, act Macros displayed high probability of interacting via ANXA1-FPR1. The interaction via CD40-CD40LG axis was predicted in several myeloid and T/NK subset combinations, but it was highest between actDC and T follicular helper (TFH), naïve and transitional CD4 T-cells. Finally, the common predicted interactions of cDC1, actDC, and act Macro clusters in TC included: IFNG-IFNGR1/2 with NK and EM CD8 T cells, CD86-CTLA4 with Tregs, exhausted CD8 T-cells, and TFH cells.

## DISCUSSION

HPV-driven TC is a subset of HNC characterized by type 1 T-cell inflammatory responses (40). Effective cellular immune responses are complex phenomena involving an interplay of different cell types. Myeloid cells represent a particularly interesting entity due to their marked ability to prime T-cells, and their wide diversity (30). Motivated by the expansion of the CD13^+^HLA-DR^+^ population in HPV^+^ TC compared to paired HT, we sought to unbiasedly assess their heterogeneity. Here, we present a high-resolution characterization of TC infiltrating and lymphoid tissue resident myeloid cells. We generated a scRNA-seq dataset of 9,505 CD45^+^CD13^+^HLA-DR^+^ myeloid cells from TC biopsies and paired HT from treatment naïve patients, using 10X Genomics. This dataset represents a tool for understanding myeloid identity dynamics in the tumor as well as steady state lymphoid tissue settings. By combining transcriptome profiling, gene signature scoring, regulon activity assessment, and RNA velocity we obtained a dynamic snapshot that illustrates the interrelationship of different myeloid communities. In addition, pathway enrichment, R-L, and survival analyses of inhouse and publicly available HNC datasets, yielded information on how myeloid communities impact survival, and the molecular mechanisms and cell-cell interactions underlying.

DC identity has recently been re-assessed in blood (30), lymphoid organs (32, 33), and several types of cancer (11, 28, 35). Here we identified 7 DC populations corresponding to 4 different lineages and diverse cell states. We observed that the cDC1 cluster was consistently homogeneous through tissue types, and its identity was dictated by the IRF8 and STAT2 regulons, as described previously (48, 49). Interestingly, cDC1s also featured high regulon activity and expression of *SMARCA5*. This TF has recently been characterized in viral DNA sensing and induction of IFN pathways, enabling DCs to prime CTL responses with a marked IFN-γ and IL-12 production (50, 51). In this context, our inhouse generated cDC1 signature highlights a positive impact of cDC1s on HNC patient survival, and TC patients when assessed separately. Pathway enrichment analysis evidenced a strong expression of genes related to MHC-I mediated cross-presentation, which is a well characterized trait of the cDC1 lineage (16). Complementing the hypothesis that cDC1s migrate in the tumor through the XCL1/2 – XCR1 axes (15), we show that besides NK cells, *XCL1*/*XCL2* were also expressed by a subpopulation of *ENTPD1+ ITGAE+* CD8 T-cells. In fact, the interaction strength between these CD8 T-cells and cDC1s was also mediated by the CLEC2B-KLRB1 axis. All in all, our data suggest a key interaction of cDC1 and a small subpopulation of TRM CD8 T-cells that has been reported to a exert tumor-reactive function in the TME of several cancer types including HNC, but also ovarian, lung, and colon cancer (52). Interestingly, RNA velocity analysis indicates that the cDC1 population stems from a cycling progenitor DC population to later mature into CCR7^+^ LAMP3^+^ actDC, suggesting that cDC1s acquire migratory potential in the TME. By comparing the signature scores of the progenitor DC population in TC and HT, we observed a marked increase in cell cycle genes and a preferential blood cDC1 signature enrichment in TC’s progenitor DCs. This indicates that progenitor DCs infiltrating TC are mainly pre-committed to a cDC lineage as is the case in blood (53). In addition, it further suggests that in the context of virally driven TC, there is a higher production rate of cDC, and that these are pre-committed to cDC1 in detriment of cDC2. This hypothesis is supported by the overall increase in myeloid APC frequency in TC, including cDC1s, while cDC2 populations were similarly abundant, as estimated both in scRNA-seq analysis but also flow cytometry. Brown et al., previously reported the presence of mitotic pre-cDCs in the spleen (33). To our knowledge, our study is the first report of cycling pre-cDCs in cancer, shifting the balance towards the cDC1 lineage.

Similar to what has been reported for blood (30) and spleen (33), DC2-like cells displayed higher heterogeneity compared to cDC1s both at RNA and protein level. Our dataset recapitulates transcriptomically distinct clusters with common *CEBPB* regulon activity (54). In scRNA-seq, MAFB^+^ LC acted as a source of DC2-like cells in TC, while RUNX3^−^ cDC2s were the unique source in HT, as seen in the RNA velocity analysis. MAFB^+^ LC uniquely exhibited MAFB and RXRA regulon activity, as well as expression of CD207, CD14, CD1A and CD1C at gene and protein level. Furthermore, these featured high scores for skin LC and tissue cDC2 gene sets (35), and no enrichment in a DC3 gene-signature, indicating that these cells did not correspond to the recently characterized DC3 subset in blood (30, 55). In fact, our flow cytometry results showed that LC and DC3 are two distinct entities judging by the lack of CD163 co-expression with CD207 and CD1a. In addition, some MAFB^+^ LC exhibited high scores of G2 and M cell cycle phases, indicating that these cells had self-renewal capacity in tonsillar tissue as they do in skin (38, 56). MAFB^+^ LCs featured high RUNX3 upregulation, and downregulated the activity of the MAFB regulon on their differentiation into RUNX3^+^ cDC2s. In line with our results, previous studies have reported expression of *MAFB* and *RUNX3* during LC differentiation in mouse mucosa and skin (57–59). In contrast, RUNX3^−^ cDC2s most likely represented *bona-fide* lymphoid organ resident cDC2s, as judged by their exclusive presence in HT, absence in TC and other HNC subtypes, as well as the lack of monocyte lineage markers and regulons at RNA and protein level. We observed that cDC2s upregulated RUNX3 along with CD207 and interferon response genes *ISG15, IFI30*, and *IFITM3* in TC. We hypothesize that, in the TME, cDC2s are subjected to IFN-I dependent maturation, potentially via TLR stimulation as it occurs upon viral or poly I:C stimulation (60, 61). Consistent with our observation at RNA level, CD207^+^CD1a^−^ cDC2s featured a more mature profile at protein level as compared to their CD207^−^ counterparts. Upon maturation into actDC, DC2-like cells downregulated the expression of *RUNX3*, which is a known repressor of *CCR7* expression and cDC2 maturation (62). Collectively, RNA-velocity analysis showed that both cDC2 and LC matured into actDC gaining *CCR7* expression, as reported previously by Bennewies et al. and Reynolds et al. (14, 39). Survival analysis based on signature scoring of the TCGA dataset did not suggest a potential contribution to HNC patient survival of neither of the three DC2-like cell populations. However, DC2-like cells, as well as cDC1s, converged into an actDC gene-program governed by *RELB* (63), and *IRF1*/*2* (64), which was associated with an increase in HNC patient survival, as assessed by gene-signature scoring. Pathway enrichment analysis suggested that actDCs perform antigen presentation via MHC-I and II in the TME and interact with diverse T-cell populations. Interaction through the CCL19-CCR7 axis is indicative of an active T-cell recruitment into TC’s TME (65, 66), while the CD40-CD40LG axis suggests a positive feedback-loop between type-1 T-cell polarization and DC maturation (67).

We also detected a rare *SIGLEC6, ILR3A* expressing DC cluster, in close transcriptomic proximity to DC2-like cells. As reported previously, DC5 identity was linked to TCF4 and RUNX2 regulon activity (33, 68). DC5s have been reported to differentiate into cDC2-like cells *in vitro* (30). Despite detecting CD11c expression on some TC derived DC5s by FC, we did not observe directionality towards DC2-like cells in RNA-velocity analysis. Together these results suggest a limited polarization into cDC2 by the TME. Nonetheless, DC5s featured higher HLA-DR, CD40 and CCR7 protein expression compared to plasmacytoid DCs. Hence, the specific functions of this rare subpopulation remain to be determined. Additionally, we detected a granulocyte lineage within the CD13^+^HLA-DR^+^ population, corresponding to mast cells with moderate expression of MHC-II related transcripts. Some studies have reported that granulocytes gain MHC-II expression during inflammation, highlighting their plasticity within the appropriate niche (69). Specifically, mast cells gain MHC-II expression upon exposure to IFN-γ, IL-4 and GM-CSF *in vitro*, although their specific ability to present antigens and activate T-cells *in vivo* is under debate (70).

We have previously demonstrated the expansion of the Mono-mac lineage in HPV^+^ TC compared to paired HT (71), which indicates an active recruitment of monocytes during carcinogenesis. In this study, we characterized a parallel differentiation process that stemmed from different monocyte polarization events, using RNA velocity. Our results challenge the *in vitro* M1/M2 paradigm, similarly to other scRNA-seq studies (11, 35). Monocytes expressing high levels of *FOS* and *FOSB* differentiated sequentially into NLRP3 and CXCL Macros. CXCL Macros were only found in latter tumor stages, featured high activity of NR3C1 regulon, and expressed high levels of *CXCL1/3/5/8*, which might mediate neutrophil recruitment in the TME via CXCR2. Indeed, several studies have reported accumulation of polymorphonuclear-myeloid derived suppressor cells in late tumor stages (72), and their association with worse prognosis in HNC patients (73). The second branch of the Mono-Mac lineage sourced from ISG Monos, which uniquely featured activity and expression of the IFNα master regulator IRF7 (74), as well as IRF8. ISG Monos expressed *TRAIL*, which may be involved in tumoricidal mechanisms, as described *in vitro* for IFNα-polarized monocytes (75). In addition, ISG Monos expressed the highest levels of *CXCL9/10/11* to potentially recruit CD8 T-cell in the TME through *CXCR3*, and displayed enrichment of TAP-independent antigen presentation via MHC-I pathway. Classically, these traits have been attributed to cDC1s (76), but they might be governed by *IRF8* in human monocytes (77, 78) as described in mice (79, 80). Interestingly ISG Monos sequentially differentiated into C1Q and act Macros, the latter of which had a moderate positive impact on HNC patient survival. Like actDCs, act Macros TF expression profile was associated with RELB and NFKB1/2, but the expression levels and gene-network activity were not as pronounced. We further validated the presence of act Macros by FC, proposing SDC2, CD163, PD-L1, and CD40 as phenotypic surface markers. Mulder et al. described similar macrophage population in lung, liver, and colon cancer; however, they linked their activity to recruitment of Treg and attenuation of T-cell responses via PDL1 and PDL2, features that we did not predict in our analysis (35).

In summary, we have provided a comprehensive characterization of myeloid cell identity and dynamics in HPV^+^ TC and HT, and we have validated our findings using 26-plex flow cytometry. Consistent with the idea that HPV^+^ TC is enriched in type-1 immune responses, we have characterized distinct polarization states of DC and Mono-Mac lineages, highlighting the impact of a type-I/II IFN-rich TME on myeloid cells. Among all populations, cDC1s stand as an ideal therapeutic target due to the biased production of their precursors, their increased abundance and capacity to mature in the tumor lesion, as well as their positive impact on TC and HNC patient survival. Our data support the conceptualization of targeted-therapeutic strategies to modulate intra-tumoral cDC1 activity.

## MATERIALS AND METHODS

The collection of tumor and contralateral tonsillar tissue was approved by the Swedish Ethical Review Authority (ref. no. 2017/580), and all participating patients granted written informed consent. Paired biopsies from TC (n=15) and contralateral HT (n=8) were obtained at Lund University Hospital at the time of diagnosis (*SI Appendix*, Table S1). The isolation of cells from human biopsies, staining and flow cytometry, scRNA-seq library preparation, sequencing, and analysis, including quality control, clustering, signature scoring, regulatory network inference, RNA-velocity, survival and RL interaction analyses, and additional datasets are described in SI Appendix, Materials and Methods.

## AUTHOR CONTRIBUTIONS

Conceptualization, D.G.J., L.G., and M.L.; Experiments, D.G.J., A.S. and C.A.; data analysis, D.G.J. and A.A.; data visualization, D.G.J.; providing human samples, S.S., D.A., and L.G.; Supervision L.G. and M.L.; Writing—original draft, D.G.J., L.G. and M.L. Writing—review and editing, all authors.

## ACKNOWLEDGEMENTS

We thank the Clinical Genomics Lund, (SciLifeLab), and Center for Translational Genomics (Lund University) for providing expertise and service with sequencing; National Bioinformatics Infrastructure Sweden (SciLifeLab) for advice in bioinformatic analysis; Cytek’s Nordic specialist team for insights in flow cytometry panel optimization; Immunology Section, Lund University for access to the Cytek Aurora instrument; and Mrs. Charlotte Cervin Hoberg, Department of ORL, Head & Neck Surgery, Skåne University Hospital, for clinical assistance. This work was supported by grants from EU Horizon 2020 Framework programme for research and innovation (EU-MSCA-COFUND, 754299 and 847583, CanFaster), the Cancera Foundation, Laryngfonden, ACTA Oto-Laryngologica Foundation, Henning and Ida Persson’s Research Foundation, Mrs. Berta Kamprad’s Cancer Foundation and the Faculty of Engineering, Lund University.

