## Supplementary information for "Single-cell analysis of myeloid cells in HPV^+^ tonsillar cancer"

**This PDF file includes:**

Materials and Methods
Supplementary References
Figures S1 to S5
Tables S1 to S3
Legends for Movie S1
Legends for Datasets S1 to S8

**Other supplementary materials for this manuscript include the following:**

Movie S1
Datasets S1 to S8

#### MATERIALS AND METHODS

##### Cell isolation

TC and HT samples were cut into small fragments in RPMI 1640 medium (Thermo Fisher Scientific, Bremen, Germany) supplemented with 0.1mg/mL gentamycin (Sigma-Aldrich, St Louis, MO). The tissue fragments were enzymatically digested with Collagenase IV (Sigma-Aldrich) (2.0 mg/mL) and DNase I (Sigma Aldrich) (200 Kunits/mL) for 20 minutes at 37° C. Cells were filtered using a 70 μm cell strainer (BD Biosciences, San Jose, CA). For scRNA-seq, $CD45^{+}$ $CD13^{+}$ HLA-$DR^{+}$ myeloid cells were isolated using FACSAria IIu (BD Biosciences) by cell sorting into FACS tubes containing 3ml of fetal calf serum (FCS) (Life Technologies, Carlsbad, CA). Sorted myeloid cells were washed twice and resuspended in 45 μL of PBS supplemented with 0.04% w/v bovine serum albumin (BSA).

##### Flow cytometry

Cells were stained with Fixable viability stain 620 (BD Biosciences) to assess cell viability. Cells were washed and non-specific binding was blocked with ChromPure mouse IgG whole molecule (Jackson ImmunoResearch, West Grove, PA) for 15 minutes at room temperature. Cells were immediately stained with the appropriate antibody panel (*SI Appendix,* Table S2 and S3) for 20 min at 4°C, washed and analyzed on a FACSAria IIu instrument (BD Biosciences) or Cytek Aurora (Cytek biosciences, Fremont, CA).

##### scRNA-seq library preparation and sequencing

The scRNA-seq libraries were prepared according to the user guide manual (CG000204) provided by 10X Genomics' Chromium Single Cell 3'Reagent Kit (v3.1) (10X Genomics, Pleasanton, CA). Briefly, sorted cellular suspensions of myeloid cells were loaded on a Single Cell chip to encapsulate single cells using the 10X Chromium controller. Encapsulated cells were then subjected to in-drop lysis and reverse transcription reactions. The emulsion droplets were disrupted, and barcoded cDNA was purified using silene magnetic dynabeads (Thermo Fisher Scientific, Waltham, MA), followed by 12-14 cycles of PCR amplification using a C1000 Touch™ Thermal Cycler with 96–Deep Well Reaction Module (#1851197, Bio-Rad. Hercules, CA). The amplified cDNA was purified with the SPRIselect reagent kit (Beckman coulter, Brea, CA) and subjected to enzymatic fragmentation, end repair, A-tailing, size selection with SPRIselect, adaptor ligation, post-ligation cleanup with SPRIselect, and sample index PCR followed by cleanup with SPRIselect. Library size and quality were assessed with High Sensitivity D1000 ScreenTape using a 4200 TapeStation System (G2991BA, Agilent, Santa Clara, CA). The cDNA libraries were sequenced on NovaSeq 6000 System (R1 – 28, i7 8, R2 – 91 cycles, Illumina, San Diego, CA).

##### scRNA-seq data analysis

Pre-processing of scRNA-seq was performed using Cell Ranger (v6.0). This pipeline included sample de-multiplexing, barcorde processing and 3' gene counting. Reads were aligned to the GRCh38 transcriptome (v. 2020-A) and single cells within each sample were merged using the aggregate function in Cell Ranger. A total of 11663 single cells with an average of 92,500 raw reads per cell were further analyzed in R (v4.0.3) and R studio (v1.4.1103) using the Seurat package (v4.0.1) (1). The datasets are available at GEO (Available upon request). A full collection of the pipeline that replicates the data analysis outlined in this manuscript is available as an R script repository on Github: Available upon request.

We used Seurat to perform thresholding, normalization, integration, dimensionality reduction, cluster identification, and visualization as well as differential gene expression analysis. The gene-barcode matrix was filtered selecting cells with more than 700 genes and less than 6000 genes detected per cell. Cells with more than 10% mitochondrial and more than 50% ribosomal UMI-counts were filtered out. Successful removal of doublets was ensured by comparing filtered cells to the barcodes selected using the DoubletFinder package (v2.0.3). Contaminating cells were identified and filtered out based on their expression of T-cell, B-cell, and NK-cell canonical genes (e.g. CD3D < 0.1 & CD3E < 0.1 & CD8A < 0.1 & CD8B < 0.1 & TIGIT < 0.1 & CD19 < 0.1 & MS4A1 < 0.1 & KLRB1 < 0.1 & CD7 < 0.1 & GNLY < 0.1 & CTLA4 < 0.1). Additionally, mitochondrial, and ribosomal genes were excluded before proceeding to data merging and normalization by log-transformation. Datasets from different samples were merged, and sample specific IDs were added to cellular barcodes allowing sample-based discrimination of cells during downstream steps. A total of 9505 single cells were further selected for sample integration, which was performed via canonical correlation analysis (CCA) using the top 2000 highly variable features across the datasets. The gene-barcode matrix was then scaled, and regression of gene expression was conducted based on UMI-counts, total number of unique UMIs, as well as the percentage of mitochondrial and ribosomal genes. Principal component Analysis (PCA) was then performed on the normalized and scaled gene-barcode matrix. The first 20 principal components were selected for non-linear dimensionality reduction (tSNE and UMAP) based on their variance using the Elbow Plot approach. The shared nearest neighbors (SNN) algorithm was used to calculate distances among cells. Cell clusters were calculated using the community detection algorithm Louvain. We inspected the granularity of the clusters using step-wise increments of 0.1 in the resolution parameter (0.1-1) implemented in the "FindClusters" function, and plotted them using the ClustTree package (v0.4.3)(2). Differential gene expression across cell clusters was performed using the "FindAllMarkers" function in Seurat. We applied Wilcoxon rank sum test to perform the analysis, only accounting for genes overexpressed in at least 10% of the cells in a cluster (p-value < 0.01, $\log_2$FC > 1).

**Gene signature analysis**

We computed signature scores using the "AddModuleScore" function in Seurat to assess cluster identity. Scores were calculated by binning signature genes in 25 bins according to average expression. The signature's average expression was corrected by subtracting the aggregated expression of 100 randomly selected genes from the same bin. We used published gene signatures for blood cDC1, cDC2, DC3, DC4, DC5, DC6, and monocytes 1-4 (3), *in vitro* activated cDC2 and moDC (4), M1/M2 *in vitro* polarized macrophages (5), skin LC (6), monocytes and monocyte-derived DC and macrophages (7, 8), and tissue mast cells (7). Additionally, enrichment scores from a pre-annotated bone marrow single cell dataset (9) were computed across our myeloid clusters in order to identify progenitor DCs. The CITE-seq bone marrow dataset was accessed through the human cell atlas (10), and a supervised PCA was computed. Supervised PCA was used to map our query myeloid dataset onto the multimodal bone marrow reference, obtaining enrichment scores for each annotation.

**Single-cell regulatory network inference and clustering analysis**

We used single-cell regulatory network inference and clustering (SCENIC) to detect and score regulons in single cells (11). The pipeline consisted of three steps: generation of a co-expression modules using Genie3, module filtering based on enrichment of DNA binding sites using RcisTarget databases, and estimation of regulon activity with AUCell. The relative activity of the top 10 enriched regulons per cluster was used to generate a heatmap. The regulon heatmap was complemented with a dot plot displaying the relative gene expression of the corresponding TF.

**Developmental pathway analysis**

We used the Velocyto python package to recount the spliced and unspliced reads. Selection of highly variable genes was performed on total reads, followed by a filtering step to select genes with more than 20 counts on both spliced and unspliced assays. After filtering, top 400-500 highly variable genes were used to calculate RNA velocity applying the scVelo "stochastic" model in the R package Velociraptor (12). Finally, we embedded the velocity vector in either a diffusion map (13) or UMAP to visualize developmental trajectories.

**Survival analysis**

TCGA-HNSC normalized gene expression data, as well as clinical metadata, were downloaded from the GDC portal repository of the cancer genome atlas (TCGA)(14). Individual samples were normalized and scored for cluster-specific gene-sets using the GSVA package (15). Then, samples were divided into quartiles, and we defined high (Q1) and low (Q4) enriched samples for each cluster-specific gene set. We performed 5 year-overall survival analysis using the survival and survminer packages (16). Difference between survival curves were assessed by log-rank test (p-value < 0.05). We used uni- and multivariate Cox proportional hazards models to estimate the relation between the gene-signature scores and the mortality rate. The assumptions of the Cox-proportional hazards model were tested assessing the Schoenfeld residuals.

**Pathway analysis**

Gene set enrichment analysis (GSEA) was performed using EnrichR (v3.0) (17) by querying differentially expressed genes among myeloid clusters onto the gene ontology database (GO, GO_bioprocess_2018)(18). We selected reoccurring and highly ranked pathways (adj.p-value < 0.01) related to APC function. Next we harmonized these pathways throughout clusters using gene set variation transformation (15). The resulting GSVA scores were plotted as a clustered heatmap using pheatmap (v1.0.12).

**RL interaction analysis**

To elucidate potential RL interactions of myeloid cells we built a "meta-dataset" containing our inhouse dataset and tumor infiltrating leukocytes from TC and HT samples of a publicly available HNC scRNA-seq dataset (19). In brief, we subsetted single cells from three TCs and five HT samples and performed normalization, sample integration via CCA, data scaling, dimensionality reduction, cluster detection and annotation based on canonical markers. Next, we integrated leukocytes of the external and inhouse datasets, normalized them via "SCTransform" and re-calculated PCA and UMAP. Once we obtained the harmonized "meta-dataset", we proceeded to RL analysis using the R package CellChat (20). First we identified RL interactions enriched in specific clusters, in TC and HT separatedly. To do so we split the "meta-dataset" into cancer and healthy, and selected overexpressed genes detected in at least 25 of the cells per cluster (p-value <

0.05, FC > 2) before calculating the communication probability. Next, we merged the cancer and healthy CellChat objects, and filtered cluster specific RL interactions that were enriched in cancer. Finally, we visualized cluster specific RL interactions of cDC1s, actDCs, and act Macros with subsets of T/NK cells using the "netVisual_bubble" function in the CellChat package.

### SUPPLEMENTARY FIGURES AND TABLES

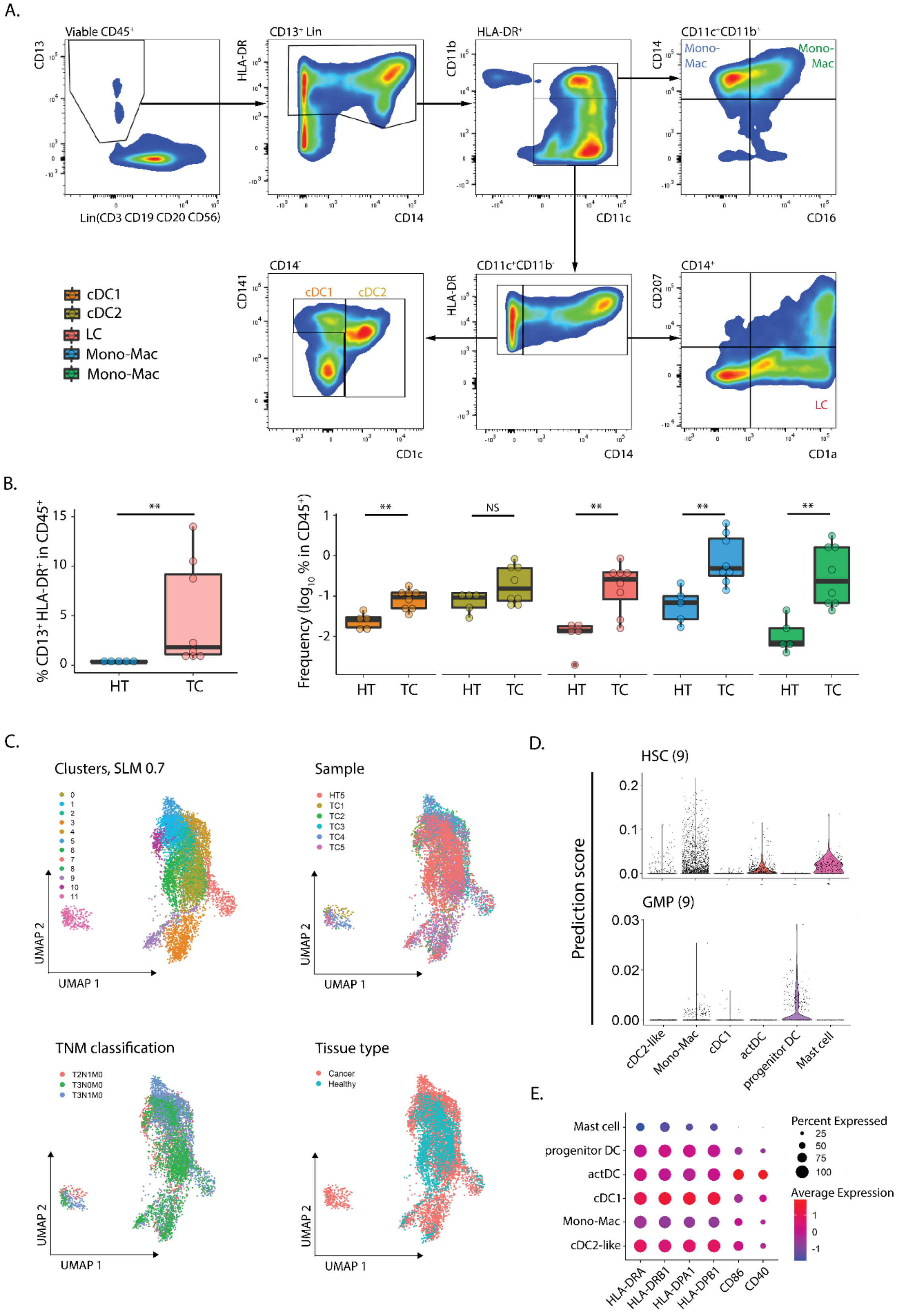

**Figure S1. Related to Figure 1.** (**A**) Gating strategy used to delineate myeloid diversity in TC and HT by flow cytometry. The following populations were identified: cDC1 ($CD11b^{-}CD14^{-}CD11c^{+}CD141^{+}$), cDC2 ($CD11b^{-}CD14^{-}CD11c^{+}CD1c^{+}$), LC ($CD11b^{-}CD14^{+}CD11c^{+}CD1a^{+}CD207^{+/-}$), and Mono-Mac ($CD11b^{-}CD14^{+}CD11c^{+}CD16^{-}$ plus $CD11b^{-}CD14^{+}CD11c^{+}CD16^{+}$). (**B**) Box plots displaying frequencies of myeloid populations identified in (A). (**C**) UMAP visualization displaying clusters calculated by SLM 0.7, sample of origin, TNM classification, and tissue type. (**D**) Prediction scores of BM hematopoietic stem-cell and granulocyte-monocyte progenitor obtained from HCA (Human Cell Atlas) (9). (**E**) Dot plot displaying expression of MHC-II related transcripts, *CD86*, and *CD40* across myeloid populations. LC: Langerhans cell; HSC: Hematopoietic stem cell; GMP: Granulocyte-monocyte progenitor.

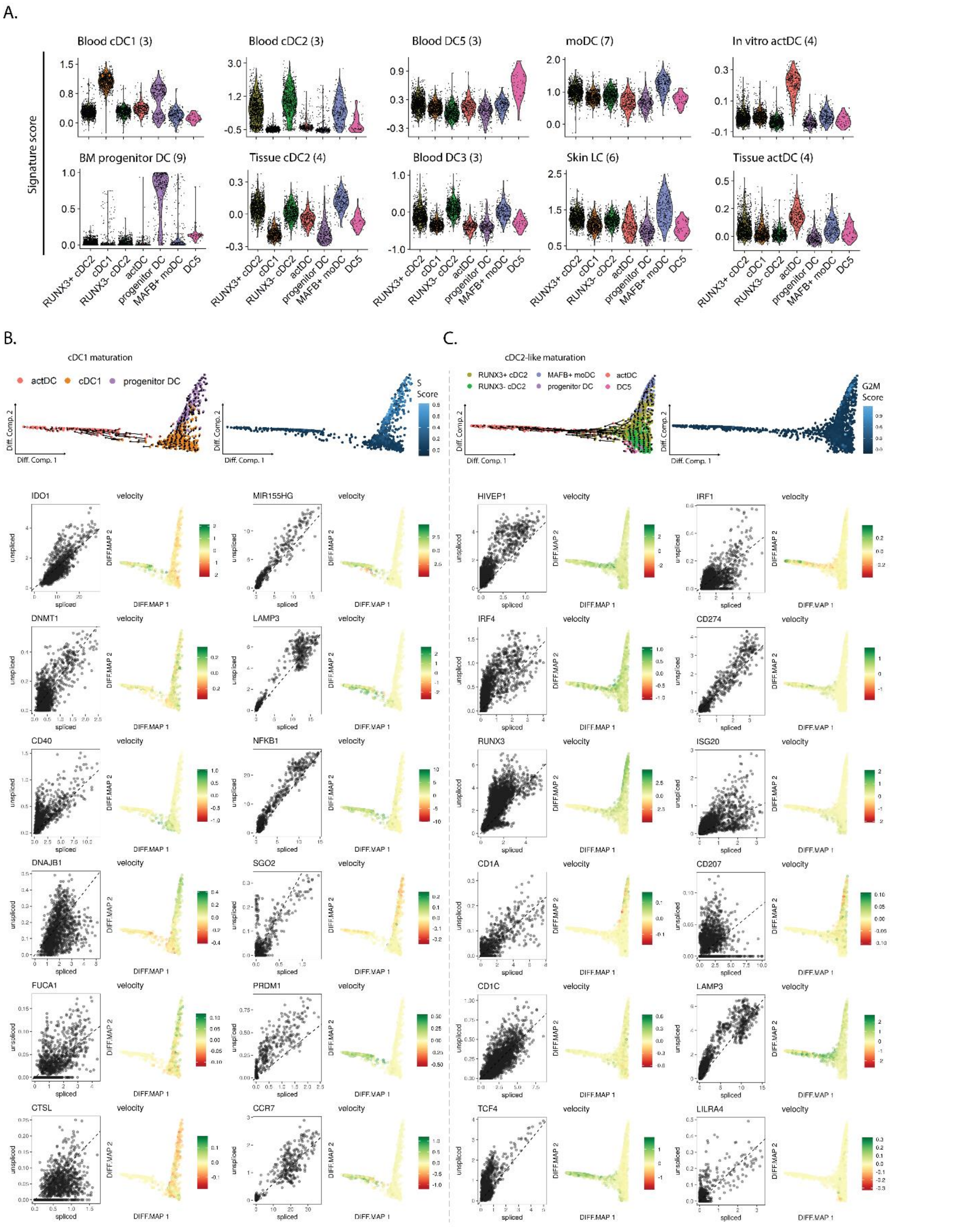


**Figure S2. Related to Figure 2**. (**A**) Gene-set enrichment scores for indicated gene-signatures. (**B**) Global and gene-specific RNA velocity and S scores in cDC1 maturation. (**C**) Global and gene-specific RNA velocity and G2M scores in cDC2-like maturation. G2M and S scores represent the relative expression of cell cycle phase genes in a cell.

A.

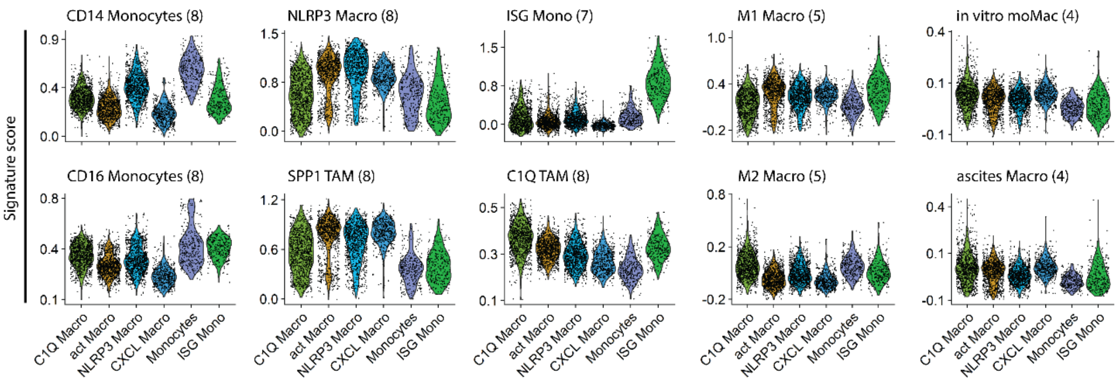


B.

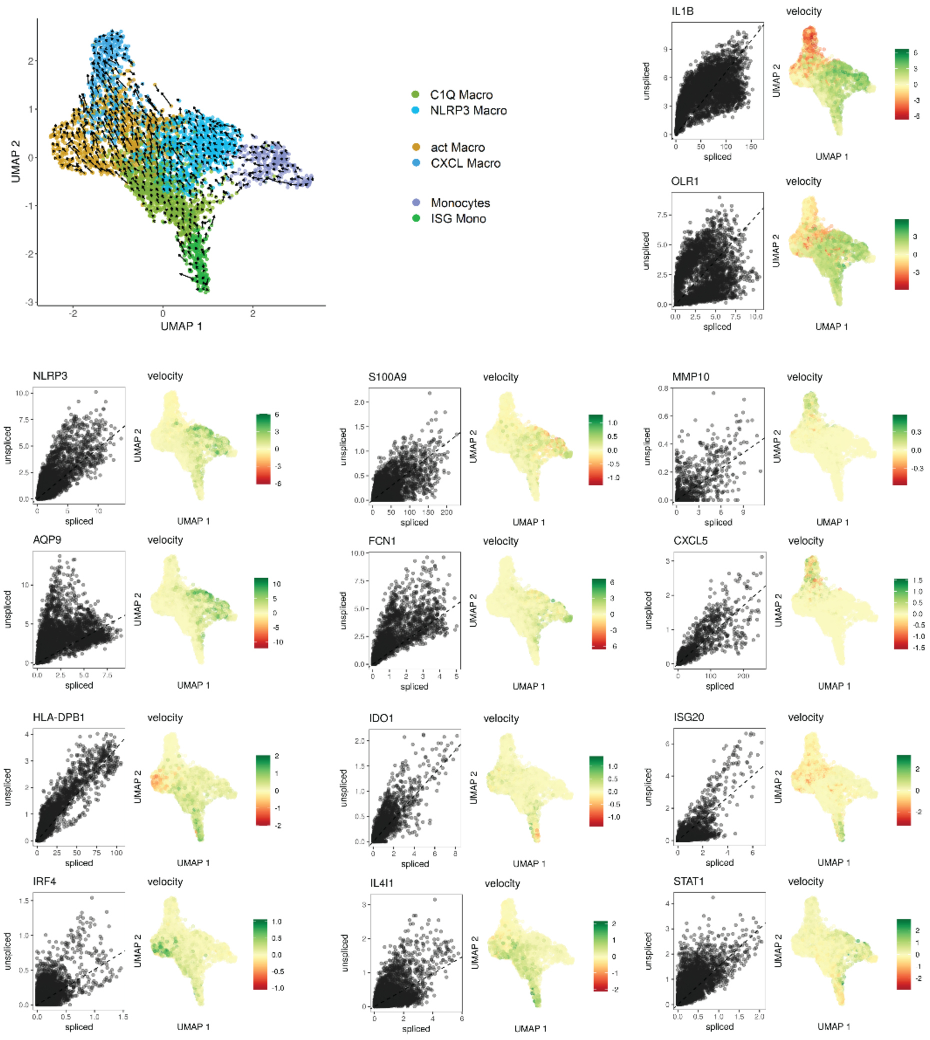


**Figure S3. Related to Figure 3.** (**A**) Gene-set enrichment scores for indicated gene-signatures. (**B**) Global and gene-specific RNA velocity of Mono-Mac lineage differentiation pathway.

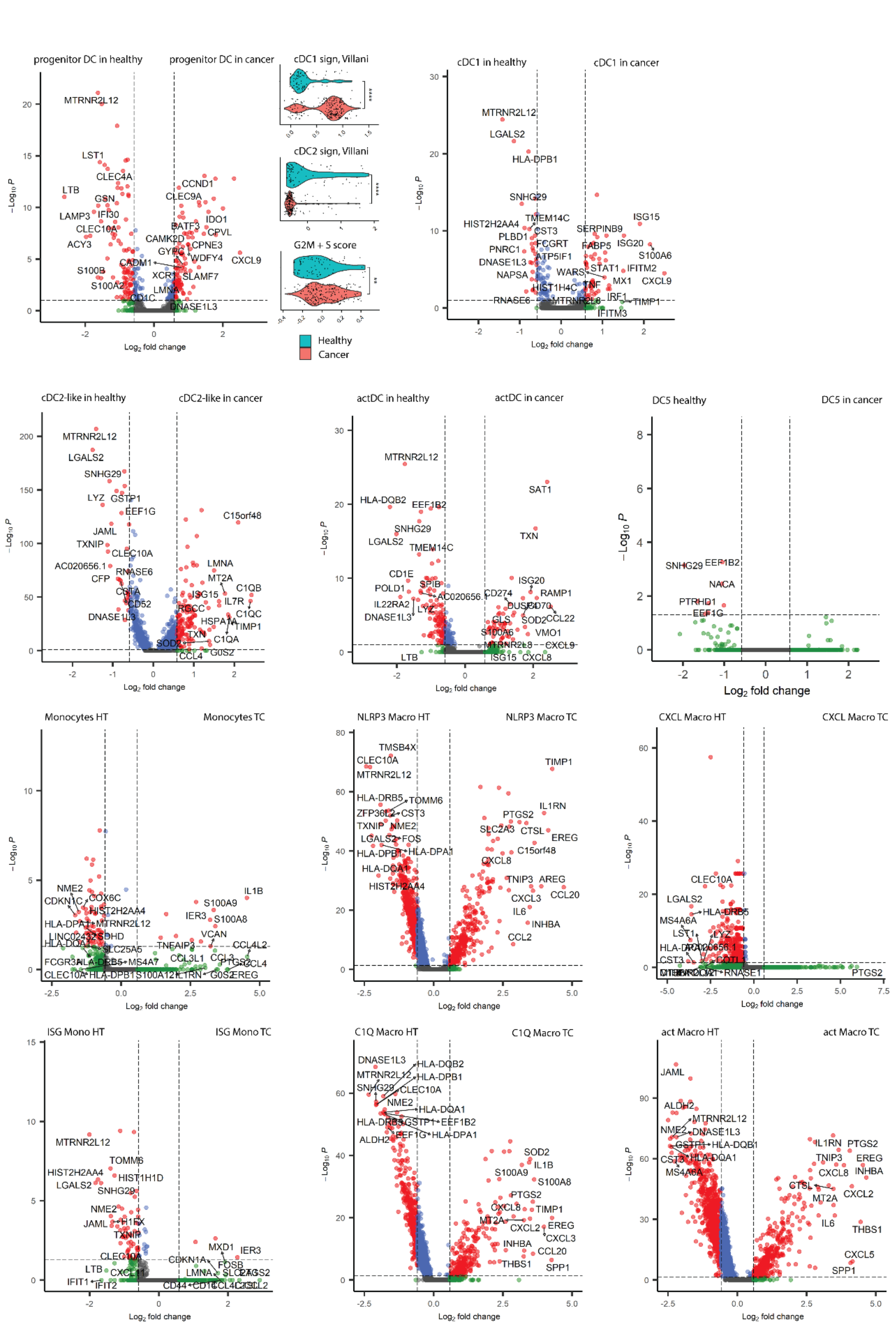


**Figure S4. Related to Figure 4.** (**A**) Volcano plots displaying DEG between TC and HT of indicated DC populations (FC > 1.5, p-value < 0.01). (**B**) Volcano plots displaying DEG between TC and HT of indicated Mono-Mac populations (FC > 1.5, p-value < 0.01). DEG: Differentially expressed genes.

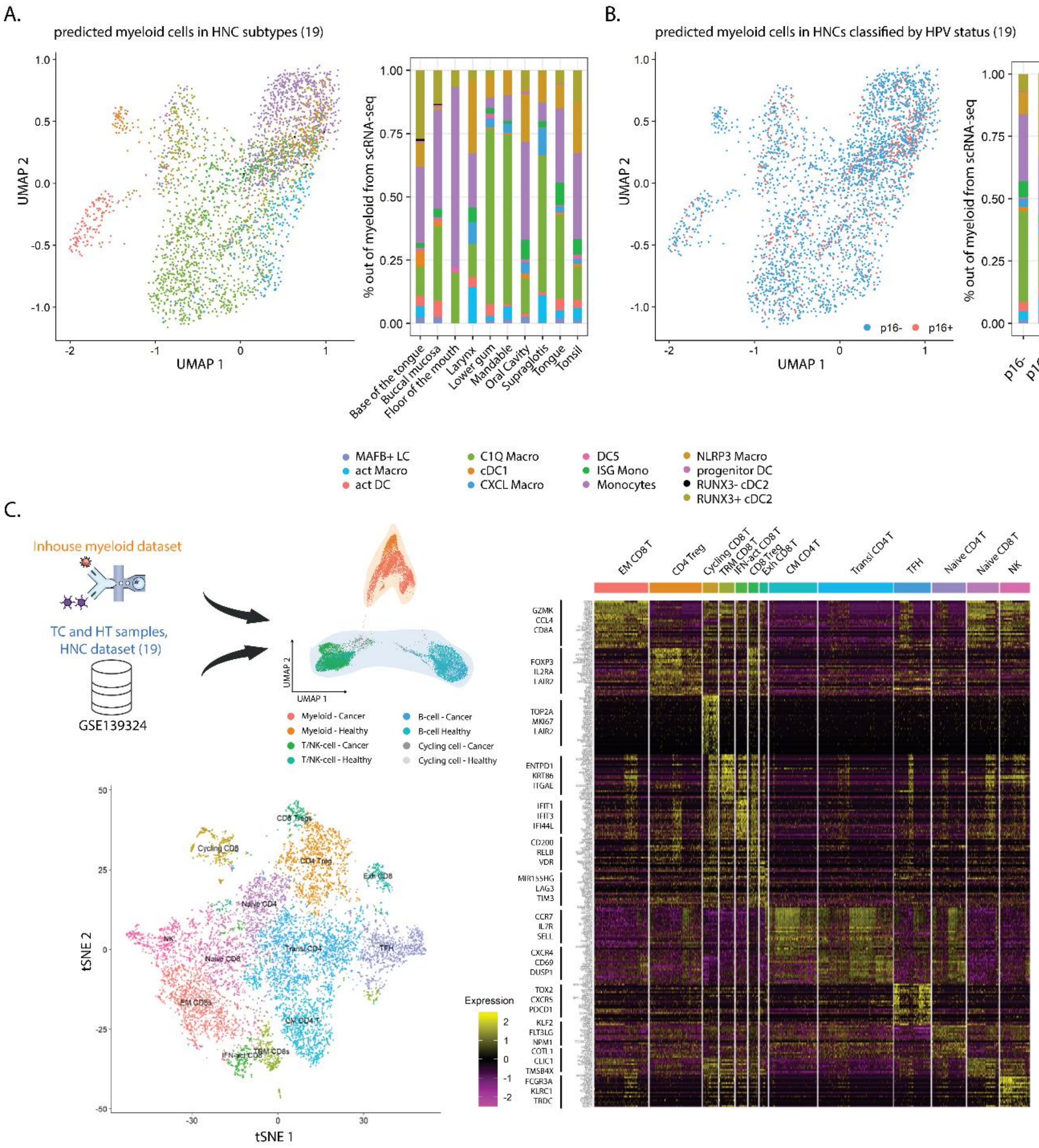


**Figure S5. Related to Figure 5.** (**A**) UMAP plot and contingency bar plot displaying myeloid populations and frequencies from a publicly available cohort of HNC (n=22) (19). (**B**) UMAP plot and contingency bar plot displaying the status of the HNC patient cohort, and myeloid frequencies based on HPV status. HPV status was assessed by the detection of p16 by immunohistochemistry. (**C**) Strategy to perform receptor ligand analysis by re-analyzing a publicly available TC scRNAseq dataset (19). UMAP displaying integrated T/NK-cell, B-cell, and inhouse myeloid lineages (top left), as well as, tSNE displaying T/NK-cell lineage subsets (bottom left) and top 25 DEG per T/NK-cell cluster (right).

**Table S1.** Patient details. FC: Flow cytometry.

| Patient ID | Sex | HPV status | TNM | Stage | Smoking status | Part of study | Matching HT |
|---|---|---|---|---|---|---|---|
| TC1 | F | P16+ | T2N1M0 | I | N | scRNA-seq and 13-plex FC | - |
| TC2 | M | P16+ | T2N1M0 | I | N | scRNA-seq and 13-plex FC | - |
| TC3 | M | P16+ | T3N1M0 | II | N | scRNA-seq and 13-plex FC | 13-plex FC |
| TC4 | M | P16+ | T3N1M0 | II | N | scRNA-seq and 13-plex FC | - |
| TC5 | M | P16+ | T3N0M0 | II | Q | scRNA-seq and 13-plex FC | scRNA-seq and 13-plex FC |
| TC6 | F | P16+ | T2N1M0 | I | Y | 13-plex FC | 13-plex FC |
| TC7 | F | P16+ | T2N0M0 | I | N | 13-plex FC | 13-plex FC |
| TC8 | M | P16+ | T2N1M0 | I | N | 13-plex FC | 13-plex FC |
| TC9 | M | P16+ | T2N2M0 | II | N | 26-plex spectral FC | 26-plex spectral FC |
| TC10 | F | P16+ | T4N1M0 | III | Q | 26-plex spectral FC | - |
| TC11 | M | P16+ | T2N1M0 | I | N | 26-plex spectral FC | - |
| TC12 | M | P16+ | T2N2M1 | IV | N | 26-plex spectral FC | 26-plex spectral FC |
| TC13 | F | P16+ | T2N1M0 | I | Q | 26-plex spectral FC | - |
| TC14 | M | P16+ | T3N0M0 | II | N | 26-plex spectral FC | 26-plex spectral FC |
| TC15 | M | P16+ | T4N2M0 | III | Q | 26-plex spectral FC | - |

**Table S2**. Antibodies used in the sorting strategy, 13-plex flow cytometry panel (**Figure 1A**; ***SI Appendix*, Figure S1A-B)**.

| **Fluorochrome** | **Antigen** | **Clone** | **Provider** | **Identifier** |
|---|---|---|---|---|
| FVS620 | Viability | - | BD | 564996 |
| APC-H7 | CD45 | 2D1 | BD | 560178 |
| APC | CD14 | Tuk4 | Miltenyi | 130-113-147 |
| PE | CD207 | DCGM4 | Beckman Coulter | IM3577 |
| PerCpCy5.5 | CD3 | UCHT1 | BD | 560835 |
| PerCpCy5.5 | CD19 | HIB19 | BD | 561295 |
| PerCpCy5.5 | CD20 | 2H7 | BD | 560736 |
| PerCpCy5.5 | CD56 | B159 | BD | 560842 |
| FITC | CD11b | ICRF44 | BioLegend | 301330 |
| BV786 | CD16 | 3G8 | BD | 563690 |
| BV421 | CD11c | B-ly6 | BD | 562561 |
| BV711 | HLA-DR | G46-6 | BD | 563696 |
| BV605 | CD141 | 1A4 | BD | 740421 |
| BV510 | CD1a | HI149 | BD | 563481 |
| PE-Cy-7 | CD13 | WM15 | BD | 561599 |
| AF700 | CD1c | L161 | Biolegend | 331530 |

**Table S3.** Antibodies used in the spectral 26-plex flow cytometry panel (**Figure 4C-F**).

| **Fluorophore** | **Antigen** | **Clone** | **Provider** | **Identifier** |
|---|---|---|---|---|
| ViaDye Red | Viability | - | Cytek Biosciences | SKUR7-60008 |
| APC-FIre810 | HLA-DR | L243 | Biolegend | 307674 |
| BV421 | XCR1 | S15046E | Biolegend | 372610 |
| cFluor B548 | CD14 | 63D3 | Cytek Biosciences | SKU R7-20116 |
| BV510 | CD1a | HI149 | BD | 563481 |
| BV785 | C1c | L161 | Biolegend | 331544 |
| PerCP Cy5.5 | CD16 | 3G8 | BD | 560717 |
| PE | CD207 | DCGM4 | Beckman coulter | IM3577 |
| APCR700 | CCR7 | 2-L1-A | BD | 566766 |
| PE-Cy7 | LAG3 | 3DS223H | Thermo fisher | 25-2239-42 |
| BV711 | CD11c | B-ly6 | BD | 563130 |
| APC | SDC2 | 305515 | RnD | FAB2965A |
| BV650 | CD11b | ICRF44 | BD | 740566 |
| BB515 | CD40 | 5C3 | BD | 565258 |
| PerCPeFluor710 | PDL1 | MIH1 | Thermo fisher | 46-5983-42 |
| PE-Cy5 | BTLA | MIH26 | Biolegend | 344506 |
| APC Vio770 | CD300E | IREM-2 | Miltenyi | 130-101-774 |
| eFluor450 | CD19 | HIB19 | Thermo fisher | 48-0199-42 |
| eFluor450 | CD20 | 2H7 | Thermo fisher | 48-0209-42 |
| PE/Dazzle594 | CD163 | GHI/61 | Biolegend | 333624 |
| BV570 | CD45 | HI30 | Biolegend | 304034 |
| cFluorR720 | CD123 | 6H6 | Cytek Biosciences | SKU R7-20013 |
| Alexa Fluor 647 | CD5 | UCHT2 | Biolegend | 300616 |
| BV480 | CCR2 | K036C2 | Biolegend | 357202 |
| BV750 | PDL2 | MIH18 | BD | 747297 |
| BV605 | TRAIL | RIK-2 | BD | 743720 |

**Movie S1 (separate file).** 3D UMAP plot displaying DC clusters, and the expression of *CLEC9A*, *CLEC10A*, and *LAMP3*.

**Dataset S1 (separate file).** DEG across DC clusters (Figure 2C).

**Dataset S2 (separate file).** Inferred regulon activity in DC clusters (Figure 2D).

**Dataset S3 (separate file).** DEG across Mono-Mac clusters (Figure 3C).

**Dataset S4 (separate file).** Inferred regulon activity in Mono-Mac clusters (Figure 3D).

**Dataset S5 (separate file).** DGEA across all myeloid clusters in TC.

**Dataset S6 (separate file).** Pairwise DGEA comparing all myeloid clusters in TC vs HT (Figure S4).

**Dataset S7 (separate file).** Pathway enrichment analysis of all myeloid clusters (Figure 5D).

**Dataset S8 (separate file).** Predicted RL interactions across all myeloid clusters, T/NK, and B cells (Figure 5E-F).
